# Partitioning variance in reproductive success, within years and across lifetimes

**DOI:** 10.1101/2022.02.08.479606

**Authors:** Robin S. Waples

**Author notes:** Corresponding author: Robin Waples.

## Abstract

Variance in reproductive success (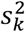, with *k*=number of offspring) plays a large role in determining the rate of genetic drift and the scope within which selection acts. Various frameworks have been proposed to parse factors that contribute to 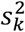, but none has focused on age-specific values of 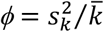, which indicate the degree to which reproductive skew is overdispersed (compared to the random Poisson expectation) among individuals of the same age and sex. Here, an ANOVA sums-of-squares framework is used to partition variance in annual and lifetime reproductive success into between-group and within-group components. For annual reproduction, the between-age effect depends on age-specific fecundity (*b*_*x*_), but relatively few empirical data are available on the within-age effect, which depends on *ϕ*_*x*_. By defining groups by age-at-death rather than age, the same ANOVA framework can be used to partition variance in lifetime reproductive success into between-group, within-group, and longevity components. Analyses of simulated data and worked examples for black bears and great tits illustrate the methods and show that the largely-neglected within-age effect a) typically represents a substantial component of the overall variance (even under a null model of random reproductive success), and b) can dominate the overall variance when *ϕ*_*x*_>1.

## Introduction

Variation among individuals is the stuff of evolution. Evolution by natural selection requires individual variation in heritable phenotypic traits affecting fitness. Over a century ago, RA Fisher (1918) introduced the term “variance” (the square of the standard deviation) as the preferred metric for measuring this variation, and immediately efforts began to identify different factors contributing to an overall variance. Perhaps the most famous such example was by Lewontin (1972), who estimated that only about 15% of the total molecular genetic variance in humans was due to differences between races or between populations within races, with the remaining 85% found among individuals within populations. The amount of data Lewontin had available at the time was extremely limited by today’s standards, but his qualitative conclusions have proved surprisingly robust and influential across a half-century (Novembre 2022).

Another kind of variance—in reproductive success, measured as the number of offspring, *k*, contributed to the next generation—is also crucially important to evolution. The ratio of variance-to-mean offspring number 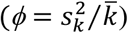has been termed the “Index of Variability” (Crow and Morton 1955), and this index is the primary factor that determines to what extent (if any) the effective population size (*N*_*e*_) is less than the census size (*N*) (Crow and Denniston 1988; Caballero 1994). A related index defined by Crow (1958; 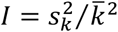 = the variance in relative fitness) has come to be known as the Opportunity for Selection because its sets an upper limit to the rate of evolutionary adaptation. Variance in offspring number appears in the numerator of both of these indices and consequently has as large role in determining both the rate of genetic drift and the scope within which selection can act.

The major goal of this paper is develop an approach analogous to Lewontin’s, but instead of apportioning genetic data based on race or geography, we will be concerned with partitioning the overall variance in reproductive success into within-age and between-age components. Emphasis is on the large fraction of the world’s species that are age-structured and iteroparous, with overlapping generations and strongly seasonal (birth-pulse) reproduction (Caswell 2001). For these species, it is important to consider two different frameworks for measuring reproductive success: within seasons or time periods (hereafter assumed to be years), and over lifetimes (quantified as lifetime reproductive success, or *LRS;* aka lifetime reproductive output, van Daalen and Caswell 2017). For annual reproduction, two systematic components contribute to the overall variance in offspring number: a between-age effect, and a within-age effect. The between-age effect depends on how expected reproductive success varies with age, as reflected in age-specific expected fecundity values (*b*_*x*_) from a life table. The within-age effect depends on age-specific values of *ϕ*_*x*_, which unfortunately are rarely reported in the literature. As a consequence, how *ϕ*_*x*_ varies across species and between ages within species is largely unknown.

For lifetime reproduction, the relevant metric is variance in *LRS*, 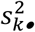(Hill 1972; Brown 1988; Tuljapurkar et al. 2020). Lifetime 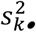is affected by age-specific variation in *b*_*x*_ and *ϕ*_*x*_, as well as another factor: longevity. All else being equal, individuals that live longer have more opportunities to reproduce, which increases disparity in lifetime offspring number between individuals and increases 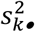. Caswell (2011) and van Daalen and Caswell (2017) use the term “Markov chains with rewards” to describe the random aspects of this process of accumulating *LRS*, and variation in longevity can be the dominant factor contributing to variance in *LRS* (e.g., Newton 1989).

The number of components of reproductive success that potentially could be identified is essentially unlimited, and a wide variety of frameworks for doing this have been proposed over the years (Arnold and Wade 1984; van Noordwijk and van Balden 1988; Brown 1988; Ferguson and Fairbairn 2001; Broekman et al. 2021). Some researchers have paid attention to within-age contributions to reproductive variance, but if so it has generally been to identify ages for which this variance is relatively large (e.g., the Siberian jay example in Engen et al. 2010; the moose example considered by Lee et al. 2020; and the sagebrush case study by Snyder et al. 2021). All of these approaches can provide useful insights, depending on one’s objectives and the kinds of data that are available, but none has focused on quantifying the overall contribution from variance in offspring number among individuals of the same age and sex. This is an important data gap; as shown below, this component of reproductive variance generally dominates the overall annual variance even when it is caused entirely by random stochasticity.

In what follows, I first describe a simple one-way ANOVA framework within which researchers can partition variance in annual reproductive success into within-age and between-age effects, and can partition variance in *LRS* into within-age, between-age, and longevity effects. Analytical methods are also developed that allow researchers to calculate what fraction of these variance components can be attributed to random stochasticity in reproduction and survival. A secondary objective is to highlight the importance of the largely neglected within-age component to the overall variance. Modern molecular tools and improved methods for parentage analysis make it easier than ever to collect empirical data on variance in offspring number. Analysis of annual data is important because any long-term study begins with data collected for individual years, and many empirical studies never encompass entire lifespans for the focal species (Nishida 1989). Furthermore, the distribution of annual reproductive success provides key insights into mating systems, and the most direct way to study sexual selection is by collecting data on all mature individuals that co-occur at the same time and place. That is not the case when analyzing only *LRS* data. Understanding the relative magnitude of between-age and within-age effects should lead to richer insights into mating systems and reproductive biology, as well as an increased ability to predict evolutionary responses to environmental changes that can affect fitness.

## Methods

See Table 1 for notation and definitions of terms.

**Table 1.** Notation

|  |  |
| --- | --- |
| $x$ | Age (in years) |
| $\alpha$ | Age at maturity |
| $\omega$ | Maximum age |
| $q$ | Age at death |
| $n$ | Number of adult age classes = number of groups in ANOVA |
| $s_x$ | Probability of surviving from age $x$ to age $x+1$ |
| $l_x$ | Cumulative probability of surviving to age $x$, with $l_x = 1$ |
| $N_1$ | Cohort size = number of offspring of one sex produced per time period that survive to age 1 |
| $N_x$ | Number of individuals of age $x$ alive at any given time; $N_x = N_1 l_x$ |
| $D_q$ | Number of individuals that died after reaching age $q$; $D_q = N_q - N_{q+1}$ |
| $N_T$ | Total number of individuals of all ages alive at any given time; $N_T = \sum N_x$ |
| $N_A$ | Number of adults with age $\geq \alpha$ alive at any given time |
| $Q$ | An index of sampling intensity relative to sampling all offspring in a stable population |
| $b_x$ | Expected number of offspring in one time period for parents of age $x$ |
| $k_{i,x}$ | Actual number of offspring produced in one time period by parent $i$ of age $x$ |
| $\bar{k}_x$ | Actual mean $k$ for parents of age $x$ |
| $\bar{k}$ | Actual mean $k$ for all parents |
| $\bar{b}$ | Parametric mean offspring number for all parents in one time period |
| $\hat{b}_x$ | An estimate of $b_x$ obtained by rescaling $\bar{k}_x$ by sampling intensity; $\hat{b}_x = \bar{k}_x / Q$ |
| $\sigma_k^2$ | Parametric variance of $k$ for all parents |
| $\sigma_{k_x}^2$ | Parametric variance of $k$ for adults of age $x$ |
| $\phi_x$ | Ratio of parametric variance to mean reproductive success in one time period for adults of age $x$; $\phi_x = \sigma_{k_x}^2 / b_x$ |
| $s_{k_x}^2$ | Unbiased sample variance of $k$ for adults of age $x$ |
| $\hat{\sigma}_{k_x, scaled}^2$ | An estimate of $\sigma_{k_x}^2$ obtained by rescaling $s_{k_x}^2$ according to Equation 5 |
| $k \bullet_{i,q}$ | Total actual lifetime number of offspring produced by individual $i$ that died at age $q$ |
| $k \bullet_q$ | Actual mean $k \bullet_{i,q}$ for all individuals that died at age $q$ |
| $s_{k_x}^2$ | Unbiased sample variance of $k \bullet_{i,q}$ for all individuals that died at age $q$ |
| $k \bullet$ | Actual mean $k \bullet_i$ for all individuals |
| $\sigma_{k \bullet}^2$ | Parametric variance in $k \bullet_i$ among all $N_1$ individuals in a cohort |
| $SSB (SSB \bullet)$ | Between-age sum of squares for annual (lifetime) reproduction |
| $SSE (SSE \bullet)$ | Error or within-age sum of squares for annual (lifetime) reproduction |
| $SST (SST \bullet)$ | Total sum of squares for annual (lifetime) reproduction |

### DEMOGRAPHIC MODEL

The focal population is isolated and iteroparous, with separate sexes. Analogous methods apply to both males and females; for simplicity data are considered for a single sex, nominally female. Reproduction follows the discrete-time, birth-pulse model (Caswell 2001), with age indexed by *x*. At age *x*, each individual produces on average *b*_*x*_ offspring and survives to age *x*+1 with probability *s*_*x*_. Age at maturity (first age with *b*_*x*_>0) occurs at age α and maximum age is ω. Newborns (age 0) do not reproduce, so *b*_*x*_ is scaled to production of offspring that survive to age 1, when they can be enumerated. Cumulative survival through age *x* is 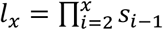, with *l*_1_=1. In a constant population with each birth cohort consisting of *N*_*1*_ yearlings, the expected number of individuals of age *x* alive at any given time is *E*(*N*_*x*_)=*N*_*1*_*l*_*x*_, and expected total census size is *E*(*N*_*T*_)=∑*E*(*N*_*x*_)=*N*_*1*_*∑ l*_*x*_. Because we are concerned with reproductive success, focus is on the adult population size, 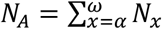.

Standard life tables provide values for age-specific survival and fecundity, to which we add a third age-specific vital rate, 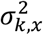, which is the variance in offspring number (*k*) around the mean for individuals of age *x* (*b*_*x*_). In many cases, it is convenient to deal with the parameter 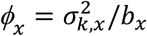, which is the age-specific ratio of variance to mean offspring number.

### VARIANCE PARTITIONING

Two major goals of this paper are to (1) quantify the relative importance of within-age and between-age effects to the overall variance in reproductive success, and (2) show how these differences depend on, and can be predicted from, age-specific vital rates. This is done using a one-way ANOVA sums-of-squares framework, which partitions sources of variation into three components (*SST*=total sum of squares of deviations, *SSB*=between-group sum of squares, and *SSE*=error or within-group sums of squares). The sums of squares are additive, such that *SST*=*SSB*+*SSE*, and the groups are the ages in an adult lifespan. In Model II (random effects) ANOVA, this partitioning commonly is done in a ‘variance components’ analysis, which directly estimates the within- and between-group variances that are associated with *SSB* and *SSE*. That approach isn’t used here, for two major reasons. First, variance-component estimation is done in a random-effects framework, whereas age is best treated as a fixed effect (Whitlock and Schluter 2014). Second, although variance components analysis is fairly straightforward in a balanced ANOVA, group sizes in a stable population (*N*_*x*_ = numbers of adults in each age class) generally decline with age. Various methods have been proposed to deal with variance component estimation in unbalanced ANOVA designs, but all have disadvantages and no consensus has emerged regarding the optimal approach (reviewed by Searle 1994).

Instead, the approach used here derives from the fact that the overall parametric variance in offspring number is 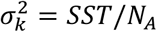, so the parametric variance partitioning becomes 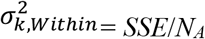and 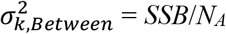, so that 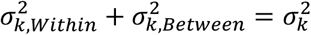. It follows that if parametric *SSE* and *SSB* can be estimated as a function of the population’s vital rates and the experimental design (including sampling intensity), it provides a means for estimating the parametric variance components. Below, this framework is used for analysis of annual and lifetime reproductive success. For simplicity in what follows it is assumed that α=1, but only minor changes to notation are required if age at maturity is >1. If α is probabilistic rather than fixed, age-specific vital rates should reflect overall means and variances across mature and immature individuals (Waples and Reed in press).

#### Annual Reproduction

Using the notation described above, the sums-of-squares components for annual reproduction are:

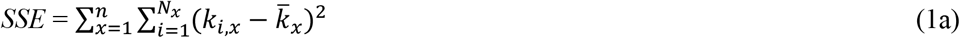

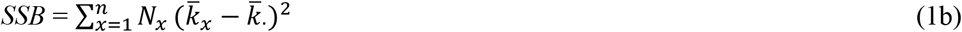

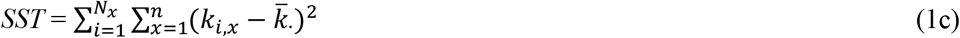

where *n* is the number of adult age classes, *k*_*i,x*_ is the number of offspring produced by parent *i* of age *x, N*_*x*_ is the group size for age class *x*, and 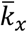and 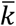are group and overall mean offspring numbers, respectively.

#### Lifetime Reproductive Success

In the lifetime reproductive success (*LRS*) version of the ANOVA framework, individuals are grouped by age-at-death (*q*) rather than age, and the symbol • is used to designate lifetime variables. For *LRS*, the sums of squares become

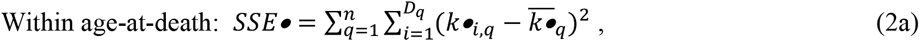

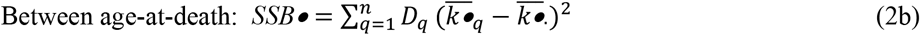

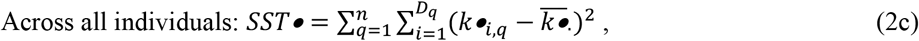

where *D*_*q*_ is the number of individuals that died after reproducing at age *q* but before reaching age *q*+1, *k*•_*i,q*_ = the total lifetime number of offspring produced by individual *i* that died at age *q*, 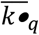 is the mean *LRS* for individuals that die at age *q*, and 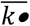 is mean *LRS* across all *N*_*1*_ individuals in a cohort. As with annual reproduction, *SST*•=*SSB*•+*SSE*•.

### SAMPLING DESIGNS

For both annual and lifetime reproduction, two sampling designs were considered. In Case 1 (comprehensive sampling), all offspring from a stable population are sampled and assigned to parents. This is the most straightforward design to evaluate because results for empirical data can be related directly to analytical expectation based on parametric vital rates. For annual reproduction, a stable population of *N*_*A*_ adults produces 2*N*_*1*_ offspring each year, so under comprehensive sampling the total number of offspring that are sampled is *N*_*Offspring*_=2*N*_*1*_, and the overall mean is 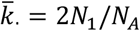. For lifetime reproduction, a stable population occurs when the *N*_*1*_ individuals in a single-sex cohort each produce an average of two offspring over their lifetime, which again means that a total of 2*N*_*1*_ offspring are sampled under Case 1. The difference is that for annual reproduction, the offspring are assigned to 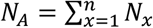potential parents while the number of potential parents in each *LRS* cohort is only *N*_*1*_, so mean offspring number per parent for annual reproduction is a fraction of what it is for *LRS*.

Case 2 (generalized sampling) includes Case 1 as a special case but allows for a variety of different experimental designs and a range of sampling intensities. Often only a subset of offspring are sampled, leading to *N*_*Offspring*_<2*N*_*1*_. Conversely, if young juveniles are sampled in a highly fecund species, *N*_*Offspring*_ can be >>2*N*_*1*_. Divergence of *N*_*Offspring*_ from the stable-population expectation leads to variation in realized mean offspring number. The ratio *Q*=*N*_*Offspring*_/(2*N*_*1*_) is a useful index of sampling intensity, with *Q*=1 replicating Case 1 (comprehensive sampling). It follows that for annual reproduction 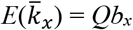. Variation in sampling intensity adds complexity to analysis of empirical data because, except in the special case of a Poisson distribution, the variance in offspring number is positively correlated with the mean (Crow and Morton 1955; Waples 2002, 2020). This in turn complicates comparisons across populations and years, and even between males and females when the sex ratio is uneven.

If *Q*≠1, it is desirable to standardize the analyses by estimating the expected variance in offspring number when the mean is the value that will produce a stable population (as suggested by Crow and Morton 1955 and many others). Using our notation for annual reproduction, the rescaling can be done using an age-specific version of the method of Crow and Morton (1955), as modified by Waples (2002):

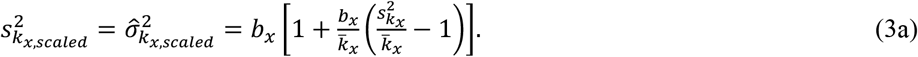

The unbiased sample variance for age *x* in the raw data is 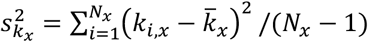, and 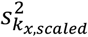is a rescaled version that represents what the variance would be expected to be if sampling had been at the level that would produce mean offspring number = *b*_*x*_, instead of the actual raw 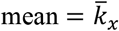. Hence, 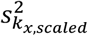is an estimate of the parametric age-specific variance in offspring number, obtained by rescaling the raw data. Since 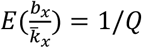, Equation 3a can be written as

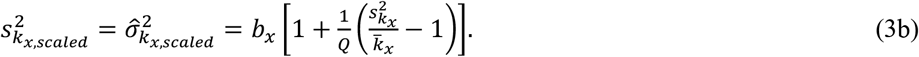

Crow and Morton (1955) derived their equation for *Q*>1, in which case Equation 3 provides the expected variance if offspring had been randomly subsampled until mean offspring number reached the target level (*b*_*x*_). Waples (2002) showed that the same formula applies when *Q*<1, in which case 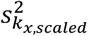is the expected result if the same offspring distribution had been sampled more intensively.

This rescaling is done separately for each age or age-at-death, but using a common *Q* based on the total number of offspring sampled. For analysis of *LRS*, the above equation is valid after replacing *x* with *q* for age-at-death, and replacing the annual metrics with their respective lifetime analogues.

The following assumptions are made regarding the estimation process:

- Means and variances in offspring number are calculated with respect to the total number of adults alive at a given time. Including juveniles, which by definition cannot produce offspring, has predictable consequences related to zero-inflation (Waples and Reed 2022) but is not considered here.
- The researcher has known or estimated ages for all potential parents in the population, so the vector *N*_*x*_ is known or can be estimated.
- Empirical data provide sample estimates of relative fecundity at age 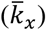 and age-specific variance in offspring number 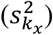, conditional on 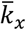.
- From the vector of relative 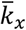estimates, the researcher can estimate the *b*_*x*_ values required to produce a stable population, using the expectation that ∑*l*_*x*_*b*_*x*_ = 2. The vector *l*_*x*_ can be estimated from observed *N*_*x*_ values, or from independent data.
- The empirical estimate of *Q* to be used in rescaling the raw data is calculated as 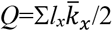.

### SIMULATIONS

Computer simulations were run to confirm the accuracy of analytical results and to evaluate performance of the estimators. Main features of the simulations are summarized here; more details and computer code can be found in Supporting Information. All simulations were done in R (R Core Team 2021) and modeled hypothetical populations with vital rates shown in Table 2. For both annual and lifetime reproduction, simulations were conducted for Case 1 and Case 2 sampling. In Case 1, the comprehensive sampling effort was fixed at *N*_*Offspring*_=2*N*_*1*_ (*Q*=1), and *N*_*1*_ was varied across a 40-fold range [50-2000] to evaluate effects of population size (and hence group size in the ANOVA analyses). For Case 2, *N*_*1*_ was set to either 100 or 500, and sampling effort was varied across an order-of-magnitude range (*Q* = [0.2,0.4,1,2]).

**Table 2.** **Top:** Vital rates for a hypothetical population having 5 age classes, constant survival (*s*_*x*_) at 50%/year, maturity at age 1, fecundity (*b*_*x*_) that is either constant, increases, or decreases with age, and variance in offspring number among individuals of the same age that is either random (*ϕ*=1) or substantially overdispersed (*ϕ*=10). *b*_*x*_ values have been scaled to values that will produce a stable population. Computer simulations used different combinations of these vital rates. **Bottom:** The range of population sizes modeled in the simulations. The vector of age-specific *N*_*x*_ values (which are also the age-class group sizes in the ANOVA analyses for annual reproduction) are determined by the relationship *N*_*x*_ = *l*_*x*_*N*_*1*_, where *l*_*x*_ is cumulative survivorship through age *x*. In parentheses after the *N*_*x*_ values are the numbers of individuals (*D*_*q*_) that die after reaching age *q* but before reaching age *q*+1. The *D*_*q*_ values are the group sizes in the ANOVA analyses of lifetime reproductive success. In this example, age at maturity is 1, so age (*x*) and age at death of adults (*q*) have the same range (1-5).

| Age ($x$) | $s_x$ | Fecundity ($b_x$) | | | $\phi$ | |
| --- | --- | --- | --- | --- | --- | --- |
|  |  | constant | increasing | decreasing | random | overdispersed |
| 1 | 0.5 | 1.03 | 0.56 | 1.24 | 1 | 10 |
| 2 | 0.5 | 1.03 | 1.12 | 0.99 | 1 | 10 |
| 3 | 0.5 | 1.03 | 1.69 | 0.74 | 1 | 10 |
| 4 | 0.5 | 1.03 | 2.25 | 0.50 | 1 | 10 |
| 5 | 0 | 1.03 | 2.81 | 0.25 | 1 | 10 |

| $x$ or $q$ | $s_x$ | $l_x$ | $N_x$ ($D_q$) | | | | | |
| --- | --- | --- | --- | --- | --- | --- | --- | --- |
| 1 | 0.5 | 1 | 50 (25) | 100 (50) | 250 (125) | 500 (250) | 1000 (500) | 2000 (1000) |
| 2 | 0.5 | 0.5 | 25 (12) | 50 (25) | 125 (62) | 250 (125) | 500 (250) | 1000 (500) |
| 3 | 0.5 | 0.25 | 13 (7) | 25 (12) | 63 (32) | 125 (62) | 250 (125) | 500 (250) |
| 4 | 0.5 | 0.125 | 6 (3) | 13 (7) | 31 (15) | 63 (32) | 125 (62) | 250 (125) |
| 5 | 0 | 0.0625 | 3 (3) | 6 (6) | 16 (16) | 31 (31) | 63 (63) | 125 (125) |

## Results

### ANNUAL REPRODUCTION

#### Parametric Variance Partitioning

To evaluate performance, it is necessary to establish what the true sums of squares are, which here are taken to be expected values in a stable population with demographics governed by parametric vital rates. These population parameters apply to a scenario in which all 2*N*_*1*_ offspring in a cohort have been sampled and assigned to parents, in which case 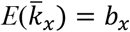 and 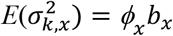 for all ages, and 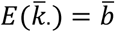, where 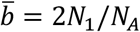is the parametric mean offspring number across adults of all ages. The parametric sums of squares expectations are obtained by substituting these terms into Equations 1a-c (see Supporting Information for details):

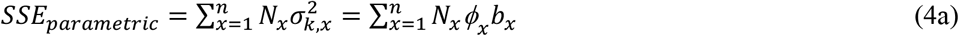

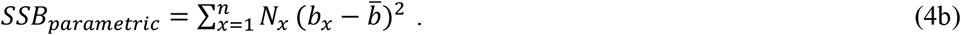

The group-size vector is *N*_*x*_=*N*_*1*_*l*_*x*_, so Equation 4a,b can be written as 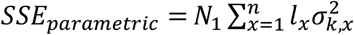 and 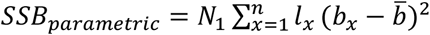. Therefore,

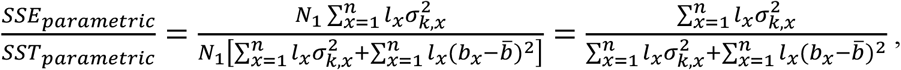

which means that the ratios *SSE*_*parametric*_/*SST*_*parametric*_ and *SSB*_*parametric*_/*SST*_*parametric*_ are independent of population size, provided that vital rates do not change with abundance.

#### A Null Model

A variety of null models might be constructed for annual reproductive success (see Waples and Reed 2022 for details), but a simple one in widespread use assumes that all potential parents function as a single Wright-Fisher population, with random mating and equal expectations of reproductive success, *E*(*k*). Under those conditions, for all ages 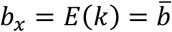and *ϕ*_*x*_= 1, which leads to

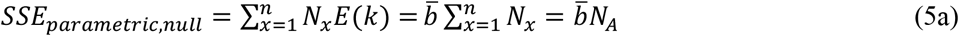

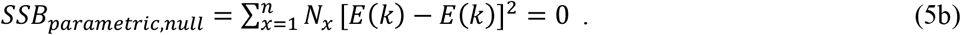

As illustrated below, this null model can provide a useful reference point for analysis of empirical data.

#### Estimation

For empirical data, Equations 4a,b are modified as follows (see Supporting Information for details):

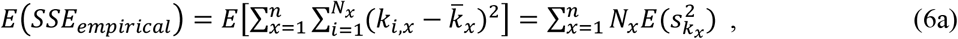

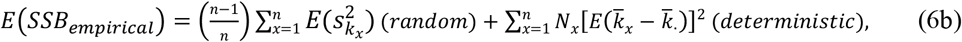

The first term in Equation 6b accounts for random contributions to 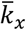, and (*n*-1)/*n* reflects the fact that the sample 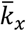values are constrained to have an overall weighted mean of 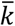, so the number of degrees of freedom is one less than the number of groups (as it is for the between-groups sum of squares in ANOVA).

Although group sizes are fixed constants, expectations for 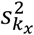 and 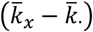depend on the sampling regime, as discussed below.

#### Case 1: Comprehensive Sampling

With comprehensive sampling, variance rescaling is not necessary, so 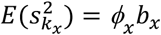and 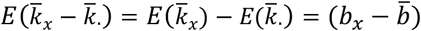and

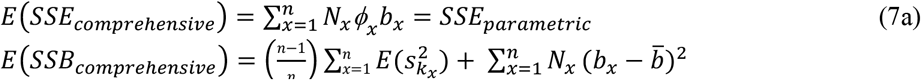

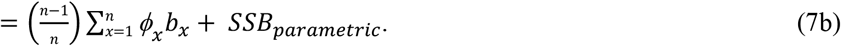

The unbiased estimator of the within-age variance 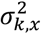is calculated from the raw data as

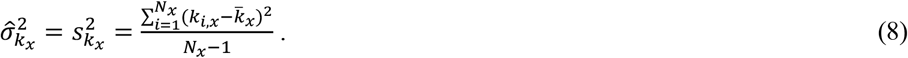

An estimator of the overall with-age sum of squares is then

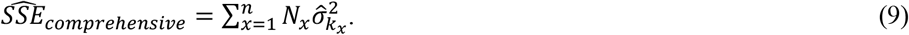

To estimate parametric *SSB* from empirical data it is necessary to subtract the expected value of the random component, leading to

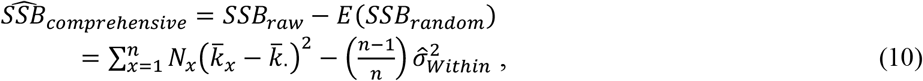

where 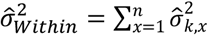.

#### Case 2: Generalized Sampling Designs

For generalized sampling, expectations for the raw, empirical sums of squares as a function of *Q* and parametric vital rates are provided in Supporting Information. The next step is to develop unbiased estimators of the parametric variance components. Estimators used in this step are:

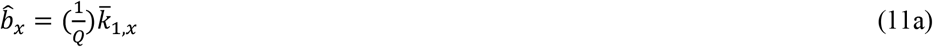

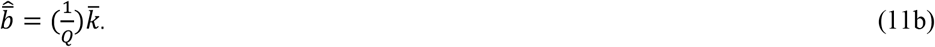

and 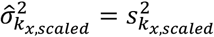is computed as in Equation 8. These estimators are then used in Equation 9 to get the overall within-group sum of squares estimator for generalized sampling:

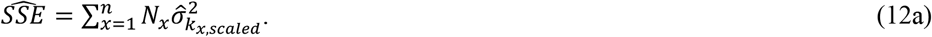

For *SSB*, it is best to account for the random contribution using the raw data and then rescale the net result to obtain the estimator of parametric *SSB* (see Supporting Information for details). The unbiased estimator of raw *SSB* is

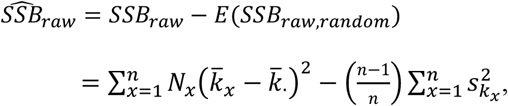

and the unbiased estimator of parametric *SSB* is

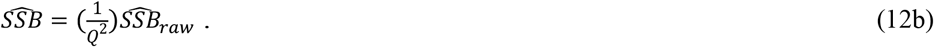

#### Simulations

Simulation results for annual reproduction are shown in Figures 1, S1-2, and Box S1. Across the scenarios evaluated, the estimators of *SSB, SSE*, and *SSE*/*SST* were asymptotically unbiased for moderate to large sampling efforts and population sizes (hence group sizes). Biases that did occur for the smaller values of *Q* (where only 20-40% of the offspring were sampled) and group size (with *N*_*1*_=50, the oldest 2 age classes have only 3 and 6 individuals) generally applied to *SSB* but not *SSE*. Because the within-age sum of squares is generally much larger than the between-age sum of squares for annual reproduction, any bias to *SSB* causes proportionally less bias to the ratio *SSE*/S*ST*, which is the primary quantity of interest. Details include the following:

- Parametric *SSE* and *SSB* both increase linearly with population/group size, but the random component to empirical *SSB* does not. As a consequence, any bias associated with adjusting for this random component becomes relatively less important as group size increases (see Box S1).
- Under a null model with no true differences in expected fecundity with age (all 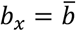), 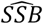was close to 0 for all scenarios, but with a slight tendency for underestimation (presumably because the correction for the random component was too large). This bias becomes smaller as sampling effort and group size increase (Figure S1).
- If fecundity increases with age (a common pattern in many species), 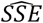remains unbiased but 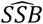slightly overestimates parametric *SSB*, leading to a slight underestimate of *SSE*/*SST*. However, the resulting biases are small even for the smallest *Q* and *N*_*1*_ (the most extreme bias occurred with *Q*=0.2 and *N*_*1*_=100, where estimated *SSE*/*SST* (0.78) was 4% higher than the parametric value, and this bias became negligible for *N*_*1*_=500; Figure 1). If fecundity decreases with age (as occurs with reproductive senescence), *SSE* is not affected but *SSB* is reduced compared to the increasing-fecundity scenario, which increases *SSE*/*SST* (Figure S2).
- For generalized sampling, raw empirical *SSE* and *SSB* (represented by filled black circles in the figures) agreed closely with expectations given by Equations S7-S8 (solid black lines). Parametric expectations under comprehensive sampling are shown in dotted red lines, which intersect the trajectories of the raw data only for *Q*=1, which represents comprehensive sampling. Red triangles show how close rescaled estimates come to these parametric expectations.
- Raw *SSE* and raw *SSB* both increase with sampling effort and hence mean offspring number, but they do so at different rates. As a consequence, the key ratio *SSE*/*SST* based on raw data can vary substantially based on sampling effort. For example, for the scenario in Figure 1 where fecundity increased with age, the proportion of the total sum of squares due to within-age effects ranged from >0.9 for *Q*=0.2 to <0.6 for *Q*=2 (black symbols and solid black lines in the bottom panels for *N*_*1*_ = both 100 and 500). All of these samples were generated by a single (albeit hypothetical) population having one parametric set of vital rates, so this result illustrates the danger of using raw, unscaled data to draw inferences about variance partitioning.
- These results for the raw empirical data indicate that any biases associated with generalized sampling designs primarily arise during the variance-rescaling process that uses the non-linear Equation 3. In this equation, 1/*Q* becomes a scaling factor that magnifies any small biases in the raw data.

**Figure 1.**
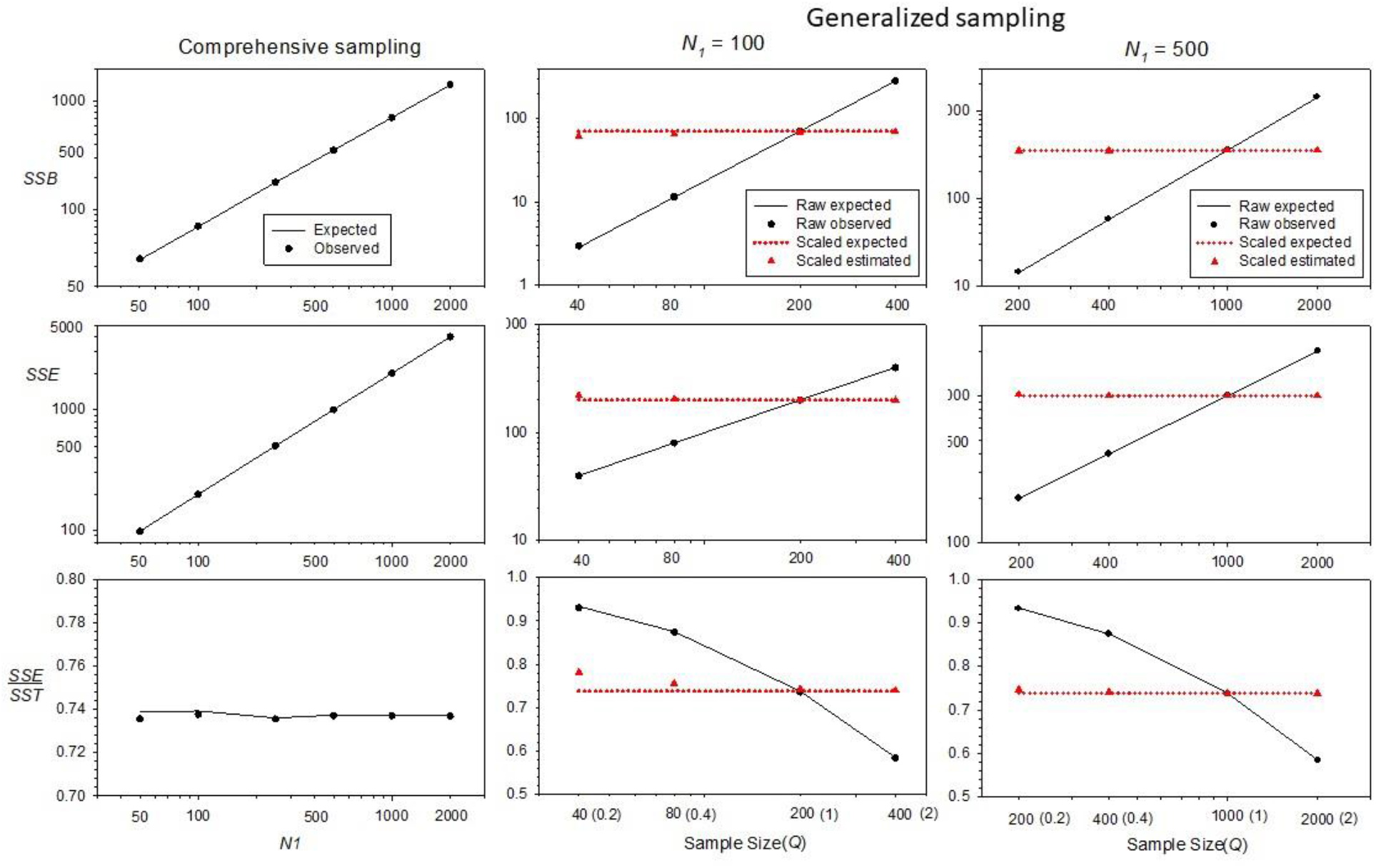
Results of simulations modeling annual reproduction in a hypothetical species for which fecundity increased linearly with age and variance in offspring number was random for individuals of the same age (all *ϕ*_*x*_ = 1; see Table 2). Left panels: Results for comprehensive sampling (Case 1) for a 40-fold range of population sizes, as indexed by the *N*_*1*_ values on the X axis. See Figure S1 for comparable results for a null model with parametric expectations given by Equations 5a,b. Black circles and black solid lines show observed and expected results, respectively. Center and right panels: Results for generalized sampling (Case 2) for four different sampling intensities, indicated on the *X* axis by *Q* = *N*_*Offspring*_/(2*N*_*1*_) = [0.2, 0.4, 1, 2]. Black circles and black solid lines show observed and expected results, respectively, using the raw data; red triangles and red dotted lines show observed and expected results after rescaling the data per Equation 3. Center panels show results for *N*_*1*_ = 100, and right panels show results for *N*_*1*_ = 500. In all cases, observed results are means across 10000 replicates.

#### A worked example—black bears from Michigan

During 2002-2010, Michigan state biologists estimated ages (from teeth) for over 2500 black bears (*Ursus americanus*) killed by hunters. This example focuses on data collected from genetic parentage analysis of a subset of bears, which yielded 221 matches of offspring to both parents (Moore et al. 2014) and allowed estimation of age-specific vital rates (Table 3). Black bears can live at least 20 years, but individuals older than 10 are uncommon, so those were grouped into a single plus age class. Males mature at age 2; females generally mature age 3 and can have litters of up to 4-6 cubs. Primary sex ratio is even, but males have lower survival so adult females are more numerous.

**Table 3.** Variance partitioning analysis for seasonal reproduction by male black bears from Michigan. Assuming constant production of *N*_*1*_ = 500 yearling males, these vital rates would produce an adult male population size of 862 age 2+ individuals. Columns under “Raw data” show estimates from field samples reported by Waples et al. (2018b). Columns under “Scaled data” show variables that have been rescaled based on the estimated index of sampling intensity *Q* = 0.05. The “Sums of squares” columns show age-specific within-age (*SSE*) and between-age (*SSB*) components. See Table 1 for notation, Table S1 for comparable data for females, and text for explanation of the calculations.

| Age | $l_x$ | $N_x$ | Raw data | | | | Scaled data | | | Sums of squares | | |
| --- | --- | --- | --- | --- | --- | --- | --- | --- | --- | --- | --- | --- |
| | | | $\bar{k}_x$ | $s_{k_x}^2$ | $\phi_x$ | $B_x$ | $\hat{b}_x$ | $\hat{\sigma}_{k_x, scaled}^2$ | $\phi_x$ | $SSE$ | $SSE_{random}$ | $SSB_{raw}$ |
| 1 | 1 | 500 | 0.000 |  |  |  |  |  |  |  |  |  |
| 2 | 0.639 | 320 | 0.027 | 0.026 | 0.957 | 8.7 | 0.54 | 0.5 | 1.0 | 172.8 | 173.6 | 0.31 |
| 3 | 0.357 | 179 | 0.030 | 0.032 | 1.066 | 5.4 | 0.60 | 1.4 | 2.3 | 248.2 | 107.3 | 0.14 |
| 4 | 0.239 | 120 | 0.030 | 0.033 | 1.101 | 3.6 | 0.60 | 1.8 | 3.0 | 216.5 | 71.8 | 0.10 |
| 5 | 0.160 | 80 | 0.062 | 0.069 | 1.121 | 4.9 | 1.23 | 4.2 | 3.4 | 335.9 | 98.3 | 0.00 |
| 6 | 0.107 | 54 | 0.121 | 0.166 | 1.370 | 6.5 | 2.42 | 20.3 | 8.4 | 1094.9 | 130.6 | 0.21 |
| 7 | 0.072 | 36 | 0.156 | 0.321 | 2.056 | 5.6 | 3.12 | 68.8 | 22.1 | 2477.3 | 112.2 | 0.35 |
| 8 | 0.048 | 24 | 0.196 | 0.329 | 1.678 | 4.7 | 3.91 | 56.9 | 14.5 | 1365.0 | 93.9 | 0.46 |
| 9 | 0.032 | 16 | 0.238 | 0.354 | 1.490 | 3.8 | 4.74 | 51.1 | 10.8 | 818.2 | 75.9 | 0.52 |
| 10 | 0.022 | 11 | 0.377 | 0.454 | 1.204 | 4.1 | 7.52 | 38.2 | 5.1 | 420.5 | 82.8 | 1.12 |
| 11+ | 0.015 | 22 | 0.122 | 0.191 | 1.560 | 2.7 | 2.44 | 29.7 | 12.2 | 653.3 | 53.7 | 0.09 |
| Totals | 2+ | 862 |  | 0.197 |  | 50 |  | 273 |  | 7803 | 1000 | 3.28 |

Cumulative survival for males though age 11 was *l*_*11*_ = 0.015 (Waples et al. 2018). Assuming a fixed number of *N*_*1*_ = 500 yearling males each year, the rest of the age classes have *N*_*x*_=*l*_*x*_\**N*_*1*_ individuals, with a total of *N*_*A*_=862 age 2+ adults. Sample estimates of age-specific fecundity increased monotonically from 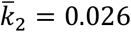for age 2 to 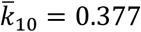for age 10, and (except for age 2) the associated sample variances 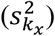 were all larger than the means, so raw *ϕ*values were >1. Expected numbers of offspring produced by each age class are 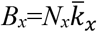, and the total across all ages is only 50, so overall sample mean offspring number is 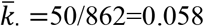. When 50 offspring is compared with the 2*N*_*1*_=1000 expected for a stable population, the index of sampling effort becomes *Q*=50/1000=0.05, indicating very sparse sampling. The identical result can be obtained by comparing 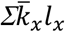 with the value expected in a stable population (2).

To estimate parametric sums of squares, the first step is to rescale the raw data to values expected for a stable population. For each age, 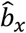and 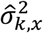are computed from Equations 11a and 12a, and the overall mean is estimated as 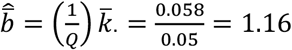. The rescaled age-specific *ϕ*_*x*_ values are considerably higher (up to 22.1 for age 7 males), indicating that overdispersion is very substantial.

These rescaled variables are then used in Equations 12a and 12b to estimate parametric sums of squares (Table 3). Overall rescaled 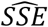is 7803. Total *SSB*_*raw*_ from the empirical data is 3.28, of which 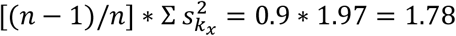can be attributed to random differences in mean age-specific fecundity. Therefore, 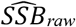 is 3.28-1.78=1.51, and 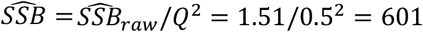. For male black bears, therefore, within-age differences among individuals are responsible for the estimated fraction 7803/(7803+601)=0.93 of the overall variance in annual offspring number. Of the total 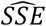, how much can be attributed to stochastic effects? Under a common null model, the distribution of offspring number within ages is Poisson, implying all *ϕ*_*x*_=1 and all 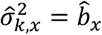, and these terms sum to 1000. So of the within-ages component, only a small fraction (∼13%) can be explained by random, Wright-Fisher reproduction. Table S1 replicates these analyses for female black bears, for which both changes in fecundity with age and age-specific *ϕ*_*x*_ are smaller than in males. The net effects on variance partitioning are similar, however, as illustrated in Figure 4.

To evaluate uncertainty associated with sparse sampling of offspring, the raw male data were bootstrapped 10000 times (see Supporting Information for details). The lower bound of the empirical 95% confidence interval (CI) for 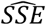was >4600, well above the null expectation (1000) if all *ϕ*_*x*_=1 (Figure 5)—indicating substantial overdispersion within ages, and the lower CI bound for 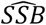was well above the value (0) expected if fecundity were constant with age. The bootstrapped CI for 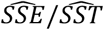was [0.804-0.966], so the point estimate that ∼90% of the total variance in annual offspring number can be attributed to variation among individuals of the same age is fairly robust, in spite of very sparse sampling.

### LIFETIME REPRODUCTIVE SUCCESS

The ANOVA sums of squares for lifetime *SSB*•, *SSE*•, and *SST*• in Equations 2a-c are superficially similar in form to Equations 1a-c for annual reproduction, but with an important difference: for analysis of *LRS*, groups are defined by age-at-death, which means that all groups with *q*>1 record cumulative *LRS* over two or more years. As shown in Supporting Information, a consequence of this is that the group-specific terms for *SSE*• take the form

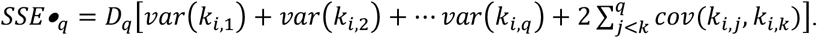

This means that within-group variances are simple additive functions of age-specific variances only if an individual’s reproduction at one age does not affect its survival or reproduction at any subsequent age. That in fact is a common assumption in modeling age-structured populations (e.g., Felsenstein 1971; Hill 1972; Waples et al. 2011), and to make the analytical expectations tractable that assumption is adopted here.

In many species, however, these covariance terms are not expected to be 0. Persistent individual differences occur when certain individuals are consistently above or below average for their age and sex at producing offspring (Lee et al. 2011), and these persistent differences lead to positive covariances in offspring production over time. Conversely, negative covariances occur when reproduction by an individual in one time period negatively affects its reproduction is a subsequent time period(s). Transient negative effects of this type are found in many species that exhibit skip or intermittent breeding (Shaw and Levin 2011; Waples and Antao 2014), and permanent negative effects can occur if reproduction adversely affects survival (e.g., McCleery et al. 1996). These temporal covariances do not affect calculation of *SSE*• from empirical data using Equation 2a, but to the extent that they do occur they will be reflected in the magnitude of the within-group sum of squares and will affect agreement with expectations based on the simpler model. Consequences of this are discussed later.

*SSB*• deals with group means rather than individuals and is not sensitive to the temporal correlations of individual reproductive success that affect *SSE*•. However, the group means 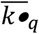are cumulative sums of *LRS* over time and hence are positively correlated. For example, 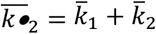and 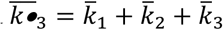share terms for mean *LRS* for individuals that die at ages 2 and 3. Furthermore, the weighted sums of squares and the weighted variance are both affected by the correlation between the patterns of change in group sample size and fecundity change with age (see Box S2 for details).

### PARAMETRIC VARIANCE COMPONENTS

As with annual reproduction, parametric values are considered to be expected values in a stable population in which all lifetime offspring have been assigned to the *N*_*1*_ potential parents in a cohort. Since the population is stable, overall mean offspring number for the cohort is 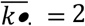.

Under those conditions, the parametric sums of squares for *LRS* are (see Supporting Information for details):

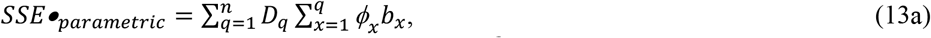

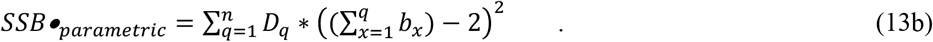

Two factors contribute to the squared-difference terms in Equation 13b: 1) changes in fecundity with age, which modulate the magnitude of 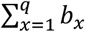, and 2) differences among individuals in age-at-death (longevity, indexed by *q*). These two factors can be separated by holding fecundity constant with age, which eliminates factor 1, so the residual *SSB*• can all be attributed to variation in longevity. If all 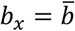, then expected *LRS* for an individual that dies at age *q* is 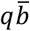, so the above equation simplifies to

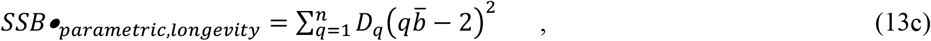

while the remainder represents the between-age component of parametric *SSB*•.

### ESTIMATION

#### Case 1: Comprehensive Sampling

Following the framework used for annual reproduction, a logical estimator of the overall with-group sum of squares for lifetime reproduction is

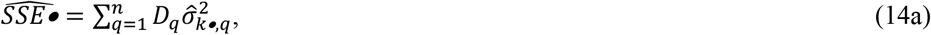

where 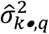 is the unbiased estimate of the variance within each group.

For comprehensive empirical data,

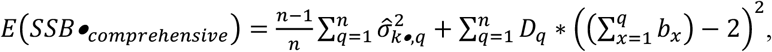

and an unbiased estimator for *SSB*• is

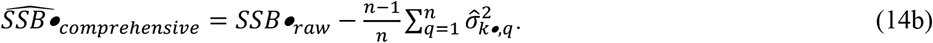

The estimator 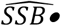 accounts for the same two factors that contribute to parametric *SSB*•: changes in fecundity with age, and differences in longevity. 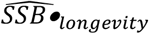can be calculated from Equation 13c using the estimator of overall mean annual offspring number 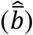 from comprehensive sampling.

#### Case 2: Generalized Sampling Designs

As before, we consider sampling at level *Q* compared to comprehensive sampling and first develop an expectation for the raw sums of squares as a function of *Q* (see Supporting Information for those results). Next we want to rescale the raw (empirical) variances to expected values under comprehensive sampling. With analogy to Equations 3 and 11a,b,

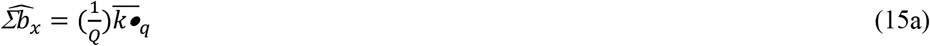

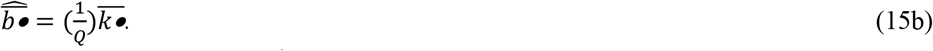

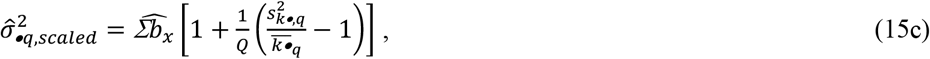

Leading to

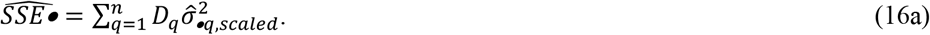

As with annual reproduction, to estimate parametric *SSB*• under generalized sampling, the first step is to adjust the raw *SSB*• to account for the random component:

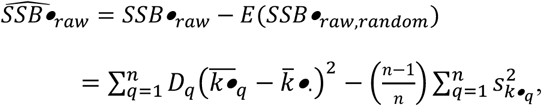

and the unbiased estimator of parametric *SSB*• is (from Equation S23)

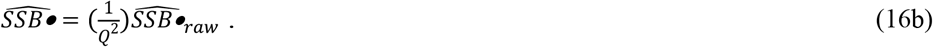

#### Simulation results

In many respects, simulation results for lifetime reproduction paralleled those for annual reproduction:

- 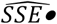 is essentially unbiased even for small group sizes and low sampling effort;
- 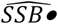 shows some minor bias for low *N*_*1*_ that largely disappears with larger group sizes;
- Except under comprehensive sampling, raw *LRS* data require rescaling to produce unbiased estimates of parametric sums of squares.

A few important differences are also worth noting. First, under the null model with constant fecundity, *SSB*• does not have an expectation of 0, as it does for annual reproduction. Individuals that die at different ages differ in number of opportunities to participate in reproduction and hence have different expectations of *LRS*. Results for *SSB*• for the null model thus can all be attributed to variation in longevity (Figures S2 and S4). All else being equal, therefore, *SSB*• makes a relatively larger contribution to overall *SST*• than annual *SSB* does to *SST*.

A second and related point is that the pattern of change (if any) in fecundity with age has a strong effect on *SSB*•. This does not lead to bias, because these effects are fully accounted for in Equations 14b and 16b. However, as shown in Figure S4 and Box S2, if fecundity declines with age (as it can with reproductive senescence), total *SSB*• can be less than would be expected if fecundity were constant (i.e., total *SSB*• < *SSB*•_*longevity*_). The interpretation in this case would be that the pattern of between-age differences in fecundity reduces overall *SSB*• compared to what it would be if *SSB*• only reflected differences in longevity.

Finally, positive or negative correlations in reproduction over time can have a strong influence on *SSE*•, whereas they have no effect on annual *SSE* because the latter considers only one time period. The example in Figure 3 simulated a population using a generalized Wright-Fisher model (Waples 2022), where individuals were allowed to have unequal probabilities of producing offspring, as indicated by a vector of parental weights, ***W*** (see Supporting Information for details). Randomly scrambling the weights each year satisfies the assumption of independence across time, producing results shown in the first half of the replicates in Figure 3. Allowing individuals to retain their weights throughout their lifetimes (second half of the replicates) creates persistent individual differences and positive correlations in individual reproductive success over time, which substantially increase *SSE*• (and hence *SST*•) but have no effect on *SSB*•.

#### Worked example – great tits

The great tit (*Parus major*) is a woodland passerine with a wide distribution in Europe and the UK. Four Dutch populations have been intensively monitored since the 1950s (Visser et al. 2021). Study sites are wooded areas fitted with an abundance of nest boxes; each year, every female that lays a clutch is netted and her ID recorded. Chicks are banded before fledging to allow tracking in the future. Data used here pertain to the cohort of birds at the Hoge-Veluwe site that matured at age 1 in 1980.

Although great tits occasionally live to 8-9 years, life expectancy is 2 years or less. In this cohort, females reproduced only at ages 1-4, so for analysis of *LRS* we consider *n*=4 groups with ages-at-death *q*=1-4. Raw data are first tabulated into a matrix with one row per female:

| ID | Age1 | Age2 | Age3 | Age4 | LRS | q |
| --- | --- | --- | --- | --- | --- | --- |
| 1 | 0 | 0 | NA | NA | 0 | 2 |
| 2 | 1 | 1 | 0 | NA | 2 | 3 |
| 3 | 0 | NA | NA | NA | 0 | 1 |
| 5 | 1 | NA | NA | NA | 1 | 1 |
| 6 | 0 | NA | 2 | 3 | 5 | 4 |

Columns 2-5 show the number of age-1recruits produced by each female at each age, with *LRS* being the total. A “0” indicates the bird was recorded attempting a nest that year but produced no recruits that were recorded in subsequent years; “NA” indicates the bird was not observed that year. Age-at-death (*q*) was taken to be the oldest age for which is.na=FALSE. In this matrix, female 1 died after age 2 without producing any recruits, female 2 produced 1 recruit at ages 1 and 2 before producing a clutch (but no surviving recruits) at age 3, and female 6 produced a clutch but no recruits at age 1, was not observed at age 2, and then produced 2 and 3 recruits, respectively, at ages 3 and 4, so its *LRS* is 5.

Females can be grouped by age-at-death to produce results shown in Table 4. The cohort includes ∑*D*_*q*_=81 females, with *D*_*1*_=48 dying after reproducing at age 1, and just 5 that reproduced at age 4. The column ‘*B*_*q*_’ shows total *LRS* for all members of each group, so 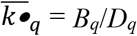 and 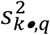is the unbiased sample variance in *LRS* for each age-at-death. The sample variance is slightly overdispersed for 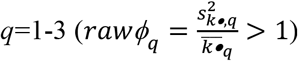, but 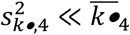, indicating substantial underdispersion in the oldest age-at-death group. This group includes only 5 individuals that, by luck or pluck, all left 3-5 total offspring. Overall mean *LRS* for the cohort is 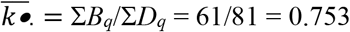, much less than mean(*LRS*) expected for a stable population (2), so *Q* = 0.753/2 = 0.376. Although birds that build nests within the study area are exhaustively sampled, reproduction also occurs in the surrounding woods, so subsequent sampling of recruits born within the study area is not comprehensive.

**Table 4.** ANOVA analysis of lifetime reproductive success in the cohort of female great tits that matured at age 1 in 1980 in the Hoge-Veluwe site in the Netherlands. The “*D*_*q*_” column shows the distribution of ages-at-death for the 81 members of the cohort. Columns under “Raw *LRS* data” show estimates of *LRS* metrics based on field samples of age-1 recruits. Columns under “Scaled data” show variables that have been rescaled based on the estimated index of sampling intensity *Q* = 0.376. For *q*=4, rescaling the raw 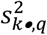 produced a negative result, so 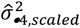was recorded as 0. The “Sums of squares” columns show within-age (*SSE*•) and between-age (*SSB*•) components. See Table 1 for notation, and text for explanation of the calculations.

| $q$ | $D_q$ | $B_q$ | Raw $LRS$ data | | | Scaled data | | | Sums of squares | | | |
| --- | --- | --- | --- | --- | --- | --- | --- | --- | --- | --- | --- | --- |
| | | | $\overline{k\bullet}_q$ | $s_{k\bullet,q}^2$ | $\phi_q$ | $\widehat{\Sigma b}_x$ | $\hat{\sigma}_{\bullet q,scaled}^2$ | $\phi_q$ | $SSE\bullet$ | $SSE\bullet_{random}$ | $SSB\bullet_{raw}$ | $SSB\bullet_{long}$ |
| 1 | 48 | 22 | 0.46 | 0.51 | 1.11 | 1.22 | 1.58 | 1.30 | 75.7 | 56.3 | 4.2 | 32.8 |
| 2 | 14 | 7 | 0.50 | 0.58 | 1.15 | 1.33 | 1.87 | 1.41 | 26.2 | 32.9 | 0.9 | 1.7 |
| 3 | 14 | 13 | 0.93 | 1.46 | 1.57 | 2.47 | 6.21 | 2.52 | 87.0 | 49.3 | 0.4 | 32.4 |
| 4 | 5 | 19 | 3.80 | 1.20 | 0.32 | 10.09 | 0.00 | 0.00 | 0.0 | 23.5 | 46.4 | 36.3 |
|  | 81 | 61 |  | 3.75 |  |  | 9.7 |  | 189 | 162 | 51.9 | 103 |

Rescaling the raw data produces estimates of parametric vital rates, based on the relationships that 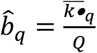and 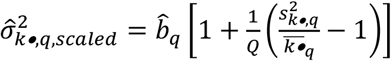. Finally, estimates of parametric sums of squares are made as follows: 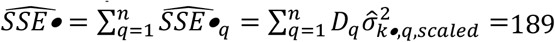. If all *ϕ*_*x*_ were 1, *E*(*SSE*•_*random*_) = 162, so most of the empirical *SSE*• can be explained by random reproduction. For the between-group sum of squares, *SSB*•_*raw*_ is 51.9, from which we subtract 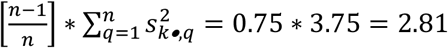to account for stochasticity, leaving 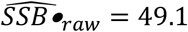. Dividing this by *Q*^2^ yields the desired estimate 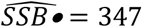. To evaluate how much of this can be explained by random variation in longevity, replace all *b*_*x*_ by 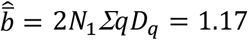. The result 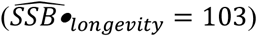 is about 30% of the total 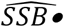, with the remainder attributed to differences in fecundity with age. The estimate of the total parametric sum of squares is 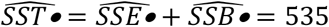, of which 189/535 = 35.3% is due to within-group effects, and most of that is attributable to random variation in reproductive success among same-age individuals (hence, overdispersion within ages is modest).

To evaluate uncertainty in these results, data for the 81 females in the cohort were bootstrapped 10000 times and the parametric sums of squares were re-estimated from each replicate. The 95% bootstrapped CI for 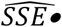 extends well below the null expectation (Figure 5, right panel), so there is no overall evidence for within-age overdispersion compared to the random Poisson expectation. The bootstrap CI for 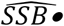 is very wide but the lower bound (105) is slightly larger than the null expectation that all between-age-at-death differences in *LRS* can be attributed entirely to variation in longevity—which is consistent with reports of some modest changes in fecundity with age in this species (Bouwhuis et al. 2012). The mean bootstrap ratio 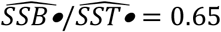agreed well with the conclusion from the original data that about two thirds of the total lifetime variance was due to *SSB*•, but the empirical 95% CI was wide (0.33-0.86), so the exact partitioning is highly uncertain.

## Discussion

Important points that emerge from results presented above can be summarized as follows:

- The ANOVA sums of squares formulas in Equations 1a-c and 2a-c do not require any assumptions about demographic or population dynamic processes and can be used with any empirical datasets that include numbers of offspring produced by each potential parent in each time period.
- Robust estimates of parametric within- and between-group sums of squares also provide robust estimates of the proportions of the total variance in offspring number arising from these two sources of variation.
- For a given age structure (relative age-group sizes determined by the cumulative survivorship vector *l*_*x*_), variance partitioning is independent of *N*, which means that randomly subsampling potential parents produces unbiased estimates of the variance partitioning (Figure S5). However, because the mean and variance of offspring number are positively correlated, variance partitioning is NOT independent of the fraction of offspring sampled. Under generalized sampling, empirical means and variances can be rescaled using an index of sampling intensity (*Q*) to allow meaningful comparisons across studies.
- Estimators of parametric sums of squares developed here are asymptotically unbiased, with modest biases to 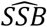and 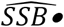 when some group sizes are <10 and/or sampling is very sparse.

Without in any way suggesting that the topic considered here is as consequential as the one Lewontin tackled in his landmark 1972 paper, some important parallels can be identified between his apportionment of human genetic diversity and partitioning of variance in offspring number. A major point of Lewontin’s paper was that the genetic differences most people were focusing on (between races, or geographic populations within races) are dwarfed by the ‘other’ ∼85% of molecular genetic variation that is found among individuals within those groups. The situation is similar for partitioning variance in offspring number, where the within-age sum of squares for annual reproduction (*SSE*) generally dominates the overall variance, even when fecundity changes sharply with age and variance within ages is Poisson (Figure S1). Published literature, however, consistently focuses primarily on the between-age component (indexed by *b*_*x*_ values from a life table) and largely ignores the within-age component. That is akin to ignoring all but the small red sectors in the black bear pie charts in Figure 4—that is, the ‘other’ 90+% of the total variation. Notably, *SSE* is also >>*SSB* (and to an even greater degree) for female black bears, which is a bit surprising, given that in most species males are expected to have higher reproductive variance. *SSE* is lower in female black bears than in males, but *SSB* is as well (and to a proportionally greater degree), leading to the net result that the within-age component is relatively more important in females.

The typical outcome of variance partitioning is somewhat different for lifetime reproduction. Mortality inevitably creates disparities in individual longevity, which increase *SSB*• and tend to make the variance partitioning more even. Still, the within-age component (*SSE*•) generally is fairly substantial and can dominate if within-age variance is overdispersed (*ϕ*>1; Figure 2).

**Figure 2.**
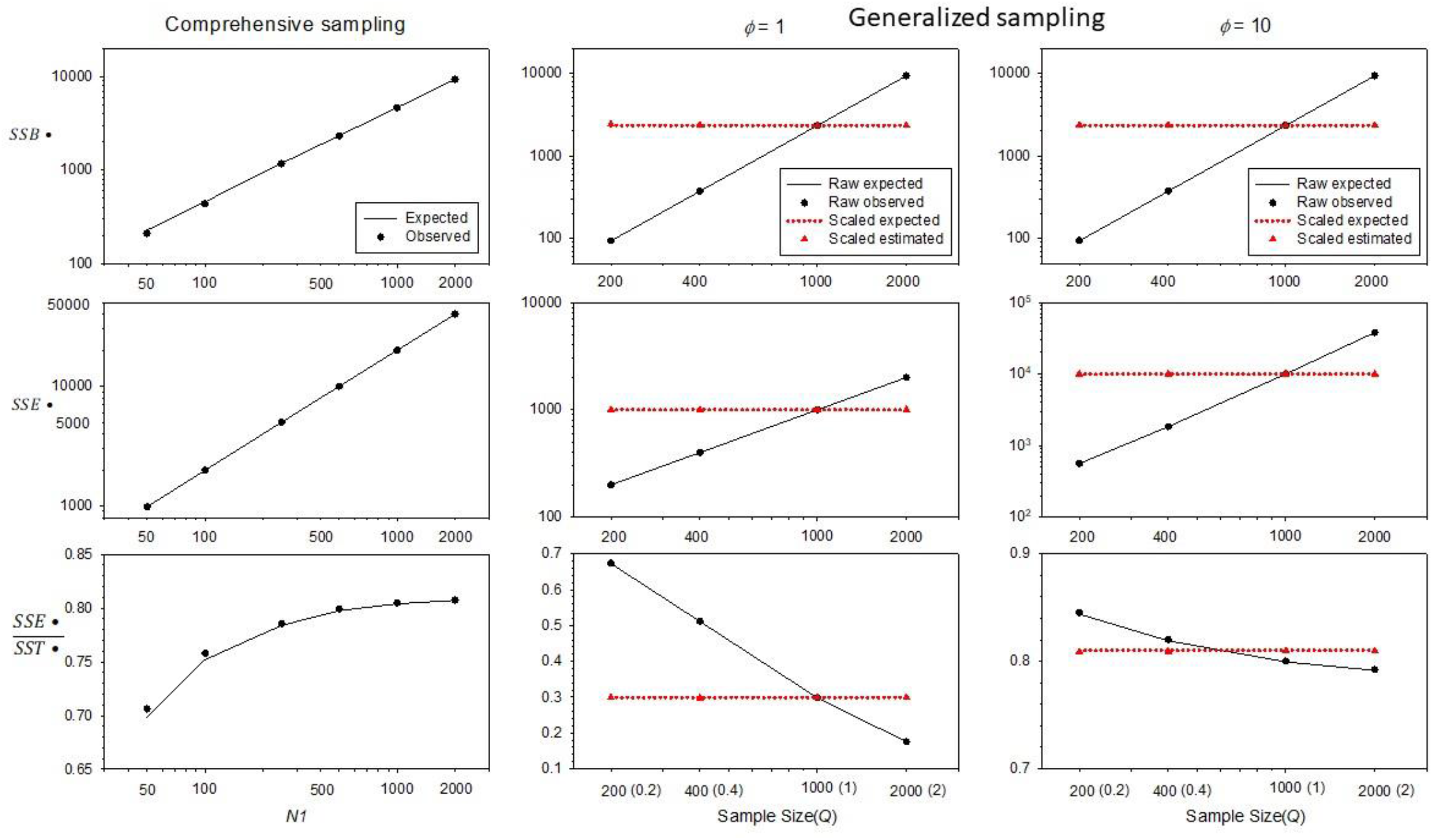
As in Figure 1, except showing results for lifetime reproductive success. For the Generalized sampling scenarios, center panels show results for *ϕ* = 1 and right panels show results for *ϕ* = 10, in both cases with *N*_*1*_ = 500.

**Figure 3.**
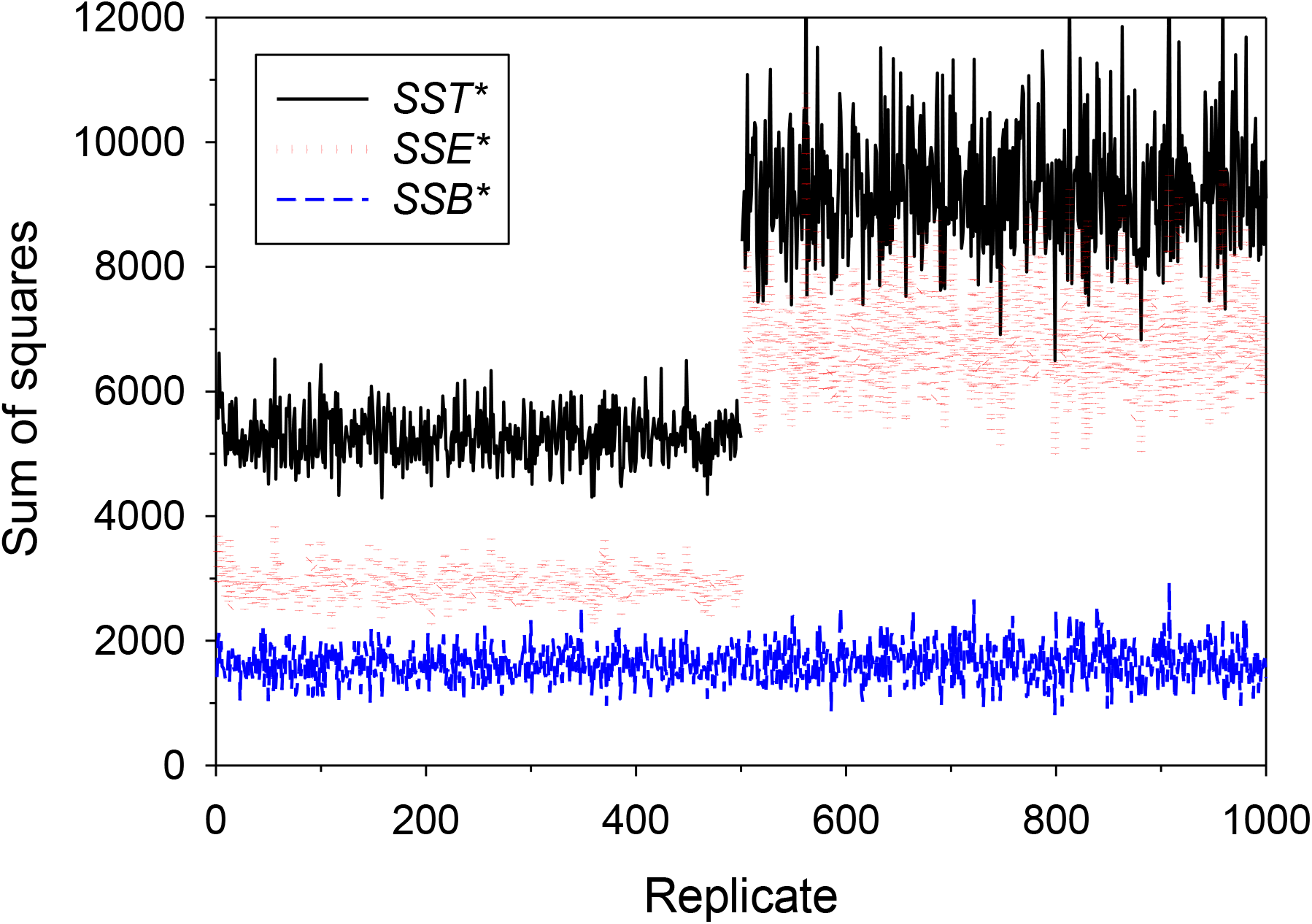
Results of simulations of *LRS* using the vital rates shown in Table 2, with fecundity that increased with age and moderately overdispersed variance in reproductive success (all *ϕ*_*x*_ = 3). The *Y* axis shows the sum of squares for the three variance components, with the groups defined by age-at-death. On the *X* axis, in replicates 1-500 parental weights were shuffled each year, so there were no persistent individual differences in reproductive success. In replicates 501-1000, the same parental weights were retained across individual lifetimes, which introduced positive correlations in realized reproductive success across time; this had no effect on *SSB* but sharply increased *SSE* and hence *SST*.

**Figure 4.**
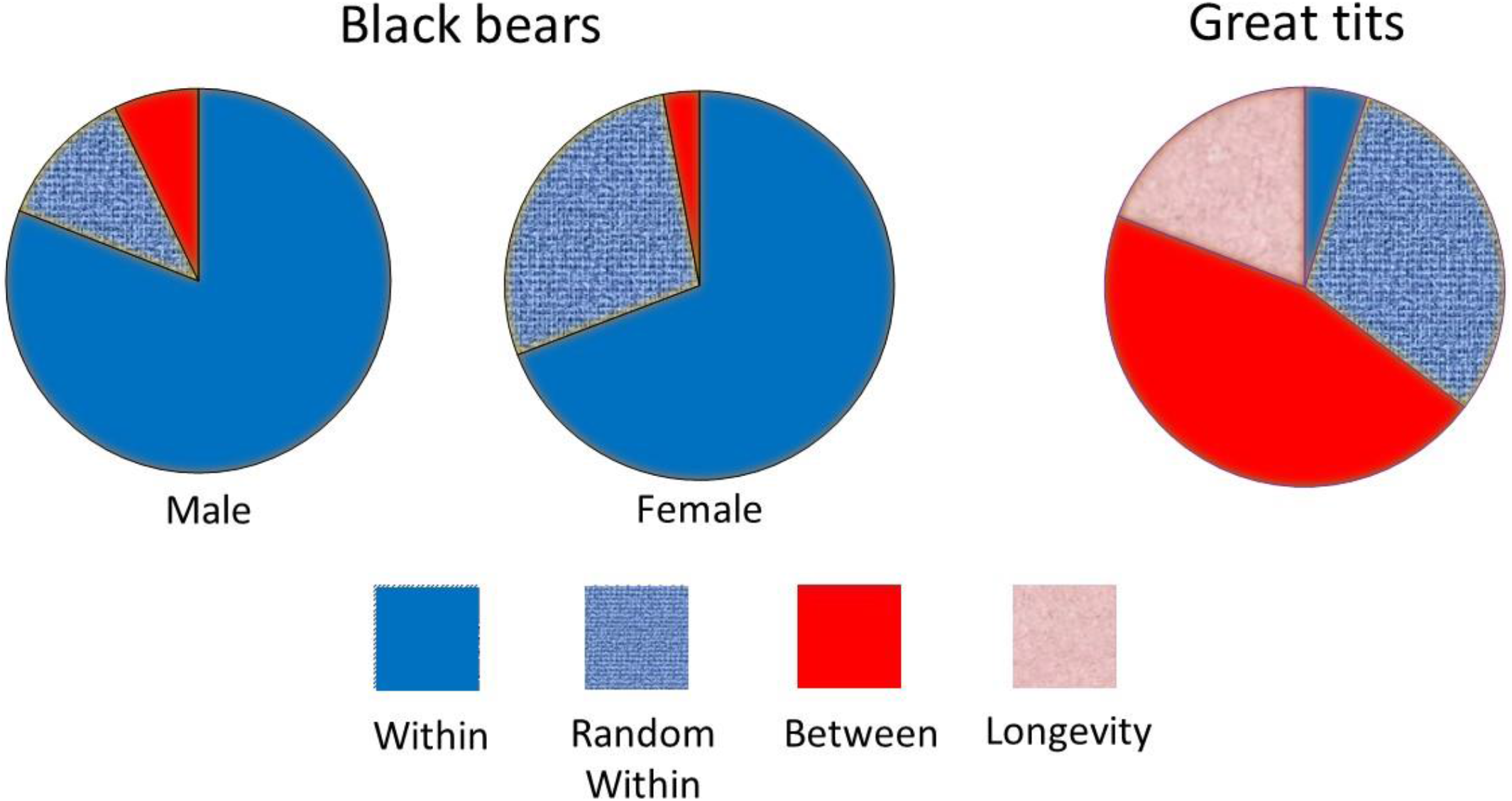
Graphical depiction of the partitioning of within-group (blue and light blue) and between-group (red and pink) components of variation in offspring number for annual reproduction in black bears (left and middle, based on data in Tables 3 and S1, respectively) and lifetime reproduction in great tits (right, based on data in Table 4).

**Figure 5.**
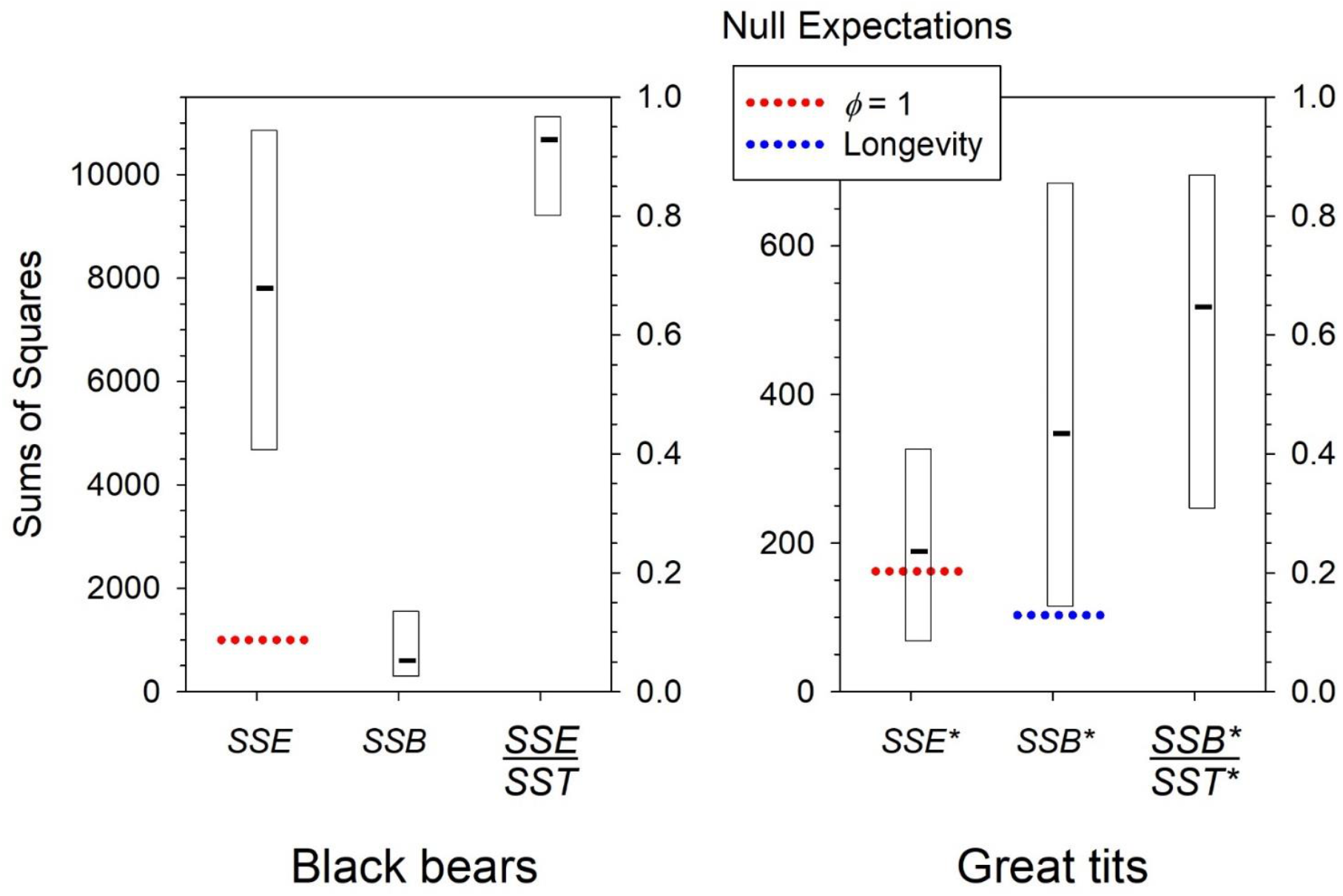
Results from bootstrapping raw data for annual reproduction in black bears (left) and lifetime reproduction in great tits (right). Black bars are point estimates discussed in the text; rectangles are empirical 95% confidence intervals across 10000 bootstrap replicates. Left axis shows sums of squares; right axis shows the partitioning of *SSE* or *SSB*• as a fraction of the total sum of squares. Dotted lines show expectations under the null model for both *SSE* or *SSE*• (red, assuming *ϕ*=1 for each age) and for *SSB*• (blue, assuming no changes in fecundity with age, in which case empirical *SSB*• would reflect only variation in longevity).

The two worked examples illustrate some of the vagaries of dealing with empirical data for natural populations. Although the black bear data were collected during an intensive study that lasted most of a decade, this represents less than half of the maximum lifespan for the species, so analysis of *LRS* was not feasible. The 221 parent-offspring matches also represented a small fraction of the estimated numbers of potential parents that might have produced matches, so effective sampling effort was very sparse (estimated at *Q*=5% for males). Nevertheless, this sparse sampling was sufficient to demonstrate with considerable certainty that, in both sexes, within-age effects account for at least 80% of overall variance in annual offspring number. Because the experimental design required combining estimates for reproduction in different years, the estimated *SSE* component includes a year effect of unknown magnitude. With more extensive data, one could estimate and account for this year effect (as done for example by Engen et al. 2005, 2010, who treated it as a random environmental effect).

Somewhat ironically, although breeding pairs of great tits are exhaustively sampled each year within the study area (as are eggs and fledglings they produce), the variance partitioning had a much higher degree of uncertainty than was found for the sparsely-sampled bears. Two factors are primarily responsible for this result. First, surviving birds can return to breed in the surrounding woods, so sampling of offspring at the recruit (age 1) stage is not exhaustive (estimated here at *Q*=37% for the cohort in question). Second, the population is relatively small, in the sampled cohort only 5 females lived to reproduce at age 4, and these birds all had high and nearly identical *LRS*. These five individuals were highly influential to the variance partitioning, as reflected in the wide range of bootstrapped results.

*LRS* data can be more complicated to interpret because of potential correlations between reproduction and survival that can affect *SSE*•. In the great tit example, 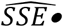 was only slightly higher than (and statistically consistent with) a null model in which within-age reproductive variance was Poisson and expected values of these correlations were 0 (Figure 5). It seems likely, therefore, that factors such as persistent individual differences and effects of reproduction on survival were not substantial, at least for this particular cohort.

The two worked examples both analyzed raw data, but the approach outlined here also can easily be used with published vital rates commonly found in life tables, provided that a third key age-specific parameter (*ϕ*) can be estimated. If the vital rates are considered to be population parameters, the parametric sums of squares can be calculated directly using Equations 4a,b for annual reproduction. Alternatively, reproduction and offspring sampling could be simulated to explore the range of results that can be expected under different experimental designs. Extrapolating from annual vital rates to *LRS* requires some assumptions; one null model was explored here, but others could be considered. In addition, for some species it might be possible to quantify temporal correlations in reproduction that can arise from different life histories. For example, the consequences of skip breeding might be modeled using the parameter *θ*_*i*_ (Shaw and Levin 2013), which gives the probability that an individual will reproduce in the current year, given that it last reproduced *i* years previously. Persistent individual differences can be modeled using *THEWEIGHT* algorithm (Waples 2022), which could be tuned to produce a desired level of correlation in individual reproductive success over time.

It has not escaped our notice that this variance partitioning approach can be used to assess the relative importance of different factors that reduce effective size compared to census size. Doing so will be the subject of a forthcoming study.

## Data availability statement

Most of the results presented here are from simulations, code for which is available in Supporting Information. Data used in the black bear example were published in Waples et al. (2018). Data used in the great tit example were provided by Marcel Visser, Netherlands Institute of Ecology, Wageningen, The Netherlands, and will be posted on acceptance. The author declares no conflict of interest.

## Notes

### Competing Interest Statement

The authors have declared no competing interest.

### Summary of Updates

Substantial rewriting throughout, but general results are unchanged

