## Supplemental info for "Partitioning variance in reproductive success, within years and across lifetimes"

### CONTENTS

Detailed Methods
Detailed Results
Tables
Figures
Box S1. Random and Deterministic Components to Annual *SSB*
Box S2. Effects of age-specific fecundity on *SSB•* for lifetime reproduction
Computer code

### DETAILED METHODS

#### Simulations

Code for conducting the simulations appears at the end of this document. For annual reproduction, the algorithm went as follows:

- Select the parametric vital rates from those shown in Table 2.
- Simulations of comprehensive sampling fixed $Q$ at 1 and considered a range of $N_1$ values.
- For each $N_1$ value, many replicates were generated where the $N_A$ adults produced a total of exactly $2N_1$ offspring.
- Group sizes ($N_x$ values) were considered fixed, and for each age class the number of offspring produced by each of the $N_x$ potential parents was chosen randomly from a negative binomial distribution with the specified target mean and variance in offspring number.
- Empirical group and overall means $\bar{k}_x$ and $\bar{k}.$ were computed from the raw data.
- If sampling was comprehensive ($Q$=1), the raw data were used directly to estimate the parametric sums of squares, as described here and in the main text.
- If $Q \neq 1$, the raw data were rescaled before estimating the parametric sums of squares.
- In one scenario using comprehensive sampling, all the offspring were assigned to all adults, and then adults and their offspring were randomly subsampled at 20-80%. This modeled a scenario in which not all adults can be sampled.

For lifetime reproduction, the algorithm differed only in the following features:

- Each replicate tracked a single cohort of offspring that had reached age 1. Individual survival from age $x$ to age $x$+1 was random, but the total numbers surviving (the $N_x$ values) were fixed based on the parametric, age-specific survivals ($s_x$). Survivors had an opportunity to reproduce at age $x$+1.
- Except as noted (e.g., in simulations producing results shown in Figure 3), the *LRS* simulations assumed independence of reproduction and survival over time, so each year each surviving individual was randomly assigned a number of offspring.

For the bootstrapping simulations for the two empirical examples, the raw data were resampled with replacement 10,000 times to construct empirical confidence intervals around the point estimates. For the black bear example, the raw data were the parents who were matched to one or more offspring through genetic parentage analysis, as summarized in Table 2 in Waples et al. (2018). For example, at age 6, 18 male bears produced exactly 1 offspring, 4 produced 2 offspring each, and one produced 3 offspring. Each bootstrap sample was then analyzed as illustrated in Table 2. For computing raw vital rates, it was assumed that the total sample of potential parents of each age was as given in Table S3 in Waples et al. (2018; column $N_x*_{(sample)}$). For the great tit example, the raw data was the matrix of LRS data for each member of the cohort, as described in the main text.

For the scenario shown in Figure 3, at age 1 each of the $N_1$ members of the cohort received a random parental weight that specified the relative probability that that individual would be chosen as the parent of a given offspring. For a given mean offspring number, the expected value of $\phi$ is a simple function of the squared coefficient of variation ($CV_w^2$) of the parental weights (Waples 2020):

$$E(\phi) \approx 1 + \bar{k}CV_w^2.$$

If every potential parent has the same weight, $CV_w^2 = 0, E(\phi) \approx 1$, and we recover the Wright-Fisher model with equal opportunities for reproductive success.

As demonstrated by Waples (2022), this equation can be rearranged to identify the $CV_w^2$ that will produce the desired $\phi$:

$$CV_w^2 = \frac{\phi - 1}{\bar{k}}.$$

For any desired $CV_w^2$, there exists a Poisson distribution with mean = variance = $\lambda$ that will produce a vector of parental weights with the desired properties. We want $CV_w^2 = var(w)/[mean(w)]^2$ = $\lambda/(\lambda)^2 = 1/\lambda$. The required mean of the Poisson distribution is thus $\lambda = 1/CV_w^2$. For example, if one wants to model reproduction with $\bar{k} = 2$ and $\sigma_k^2 = 8$, so $\phi = \frac{\sigma_k^2}{\bar{k}} = 4$, then one wants a vector of parental weights such that $CV_w^2 = \frac{4-1}{2} = 3/2$. Parental weights can be randomly generated using a Poisson distribution with $\lambda$=2/3. Randomly simulating parental weights from the Poisson distribution with this parameter will, on average, produce a vector of parental weights with $CV_w^2 = 3/2$. This process can be done separately for each age class of adults to generate offspring distributions that approximate those specified by the age-specific vital rates.

**Variance rescaling**

Equation 3b in the main text shows how empirical variances were rescaled under generalized sampling:

$$s^2_{k_{x,scaled}} = \hat{\sigma}^2_{k_{x,scaled}} = b_x\left[1 + \frac{1}{Q}\left(\frac{s^2_{k_x}}{\bar{k}_x} - 1\right)\right]. \quad (3b)$$

If the raw, sample variance in offspring number is underdispersed ($\frac{s^2_{k_x}}{\bar{k}_x} < 1$) and the sampling fraction is small enough ($Q$<1), Equation 3b can produce a negative estimate for the rescaled $\hat{\sigma}^2_{k_{x,scaled}}$. The interpretation here is that there exists no parametric variance $\sigma^2_{k_x}$ that would be expected to produce the observed sample variance $s^2_{k_x}$, given randomly sampling offspring at level $Q$. This result, however, can happen by chance, especially when the number of parents is small (as occurred in the great tit example in Table 4, where only 5 birds lived to reproduce at age 4). In this event, the best option is to record the rescaled variance as 0 (indicating that there is no real evidence it is >0).

### DETAILED RESULTS

For continuity and readability, some of the results from the main text are repeated here, but other results are presented in more detail.

#### ANNUAL REPRODUCTION

##### Parametric Sums of Squares

The population parameters we are interested in apply to a scenario in which all $2N_1$ offspring in a cohort have been sampled and assigned to parents, in which case $E(\bar{k}_x) = b_x$ for all ages, and $E(\bar{k}.) = \bar{b}$, where $\bar{b}$ is the parametric mean offspring number across adult females of all ages. The parametric sums of squares expectations are obtained by substituting these parametric expectations into Equations 1a and 1b.

Consider first each of the age-specific terms in *SSE*:

$$SSE_x = \sum_{i=1}^{N_x}(k_{i,x} - b_x)^2 = N_x E[(k_{i,x} - b_x)^2].$$

Using the definition of a variance ($Var(X) = E(X^2) - [E(X)]^2$, hence $E(X^2) = Var(X) + [E(X)]^2$),

$$E[(k_{i,x} - b_x)^2] = Var(k_{i,x} - b_x) + [E(k_{i,x} - b_x)]^2 = Var(k_{i,x}) = \sigma^2_{k,x},$$

with the simplifications coming because a) $b_x$ is a constant and subtracting a constant does not affect a variance, and b) $E(k_{i,x} - b_x) = 0$ by definition. Therefore,

$$SSE_x = N_x \sigma^2_{k,x} = N_x \phi_x b_x ,$$

and the overall parametric value for *SSE* becomes

$$SSE_{parametric} = \sum_{x=1}^{n} N_x \sigma^2_{k,x} = \sum_{x=1}^{n} N_x \phi_x b_x \quad . \qquad \text{(S1)}$$

Considering now parametric *SSB*, each age-specific term is $SSB_x = N_x(b_x - \bar{b})^2$, so the overall parametric *SSB* becomes

$$SSB_{parametric} = \sum_{x=1}^{n} N_x (b_x - \bar{b})^2 \quad . \qquad \text{(S2)}$$

### Estimation

#### Case 1: Comprehensive Sampling

For the within-age estimator, see the main text. The estimator for *SSB* has to account for the fact that in empirical data, random variation in realized mean age-specific fecundity will increase realized *SSB* above the parametric expectation shown in Equation S2. For empirical data, the expected contribution to the between-group sum of squares from individuals of age *x* is

$$E(SSB_{x,comprehensive}) = E\left[N_x\left(\bar{k}_x - \bar{k}.\right)^2\right] = N_x E\left[\left(\bar{k}_x - \bar{k}.\right)^2\right].$$

Using the variance-definition approach described above, this can be written as

$$E(SSB_{x,comprehensive}) = N_x\left[s^2_{\bar{k}_x} + \left[E\left(\bar{k}_x - \bar{k}.\right)\right]^2\right],$$

where $s^2_{\bar{k}_x}$ is the sample variance of the mean offspring number for individuals of age $x$. Now the variance of a mean is the variance of the raw data divided by the sample size, so $N_x s^2_{\bar{k}_x} = N_x \frac{s^2_{k_{i,x}}}{N_x} = s^2_{k_{i,x}}$, leading to

$$E(SSB_{x,comprehensive}) = s^2_{k_{i,x}}(random) + N_x\left[E\left(\bar{k}_x - \bar{k}.\right)\right]^2(deterministic). \quad \text{(S3)}$$

Each age-specific term contributing to realized *SSB* thus has two components: a random one related to the variance of age-specific mean offspring number, and a deterministic one related to systematic difference between the age-specific mean fecundity and the overall mean. Across all ages,

$$E(SSB_{comprehensive}) = \left(\frac{n-1}{n}\right)\sum_{x=1}^{n} s^2_{k_{i,x}} + \sum_{x=1}^{n} N_x\left[E\left(\bar{k}_x - \bar{k}.\right)\right]^2, \quad \text{(S4)}$$

with the $(n\text{-}1)/n$ term reflecting the fact that the $\bar{k}_x$ values for different ages are constrained to have an overall weighted mean of $\bar{k}.$. Note that the expected value of the sample variance is $E\left(s^2_{k_{i,x}}\right) = \hat{\sigma}^2_{k,x}$ as defined in Equation 8. Since the goal is to estimate parametric *SSB* (Equation S2), it is necessary to subtract the expected value of the random component, leading to

$$\widehat{SSB}_{comprehensive} = \sum_{x-1}^{n} N_x\left(\bar{k}_x - \bar{k}.\right)^2 - \left(\frac{n-1}{n}\right)\sum_{x=1}^{n}\hat{\sigma}^2_{k,x}.$$

$$= \sum_{x-1}^{n} N_x\left(\bar{k}_x - \bar{k}.\right)^2 - \left(\frac{n-1}{n}\right)\hat{\sigma}^2_{Within}, \quad \text{(S5)}$$

where $\hat{\sigma}^2_{Within}$ is an unbiased estimate of the within-group variance.

**Case 2: Generalized Sampling Designs**

Two questions are relevant here: 1) What are expectations for the empirical variance in offspring number and the ANOVA sums of squares under generalized sampling designs? and 2) How can raw empirical estimates be adjusted to provide unbiased estimates of the variance components?

**Within-age sum of squares estimation**

To answer the first question listed above, we can calculate $E(rawSSE_x)$ under generalized sampling as a function of $Q$ and the parametric vital rates. This can be done using a reverse form of Equation 3, where the initial data are the parametric vital rates and the rescaled variance is that expected under the actual sampling design, which yields mean offspring number of $\bar{k}_x$:

$$E\left(s^2_{k_{x,raw}}\right) = \bar{k}_x\left[1 + \frac{\bar{k}_x}{b_x}\left(\frac{\sigma^2_{k,x}}{b_x} - 1\right)\right] = Qb_x\left[1 + Q\left(\phi_x - 1\right)\right]. \quad \text{(S6)}$$

It follows that

$E(rawSSE_x) = N_x Q b_x\left[1 + Q\left(\phi_x - 1\right)\right]$, and

$$E(rawSSE) = \sum_{x=1}^{n} N_x Q b_x\left[1 + Q\left(\phi_x - 1\right)\right]. \quad \text{(S7)}$$

To address the second question listed above, the empirical vital rates are rescaled to estimate the parametric vital rates, and these rescaled estimators are then used to estimate the parametric sums of squares. The rescaled vital rate estimators are (from the main text):

$$\hat{b}_x = \left(\frac{1}{Q}\right)\bar{k}_{1,x}$$

$$\hat{\bar{b}} = \left(\frac{1}{Q}\right)\bar{k}.$$

$$\hat{\sigma}^2_{k,x,scaled} = s^2_{k_{x,scaled}} = \hat{b}_x\left[1 + \frac{1}{Q}\left(\frac{s^2_{k_{x,raw}}}{\bar{k}_x} - 1\right)\right],$$

The age-specific scaled variance estimators are then used to get the overall estimator of *SSE*:

$$\widehat{SSE} = \sum_{x=1}^{n} N_x \hat{\sigma}^2_{k,x,scaled},$$

which is identical to the formula in Equation 9, except that with generalized sampling the variance terms require rescaling.

**Between-age sum of squares estimation**

Unbiased estimation of *SSB* requires accounting for the random contribution to group means, even if the true means do not differ. Under generalized sampling designs it is also necessary to rescale the sample means and variances. There are two general options for conducting these two steps: 1) adjust the raw data for the random contribution, and then rescale the result; and 2) rescale each age-specific mean and variance, and then make the adjustment based on these rescaled values. Both options were evaluated under a variety of scenarios, with the following results:

- Both methods are asymptotically unbiased for large group sizes and sampling efforts;
- For smaller $N_1$ and $Q$, Option 1 tends to underestimate *SSB* while Option 2 tends to overestimate;
- The magnitude of the bias is generally smaller for Option 1;
- The coefficients of variation across replicates are similar for both methods.

Based on these results, Option1 was used in the analyses reported here.

Following the approach above, with $Q = N_{Offspring}/2N_1$, then for empirical data $E(\bar{k}_x - \bar{k}.) = Q(b_x - \bar{b})$, so

$$E(SSB_{x,raw}) = s^2_{k_{i,x}} + N_x[Q(b_x - \bar{b})]^2$$

$$= Qb_x[1 + Q(\phi_x - 1)]random + Q^2N_x(b_x - \bar{b})^2 deterministic, \text{ and}$$

$$E(SSB_{raw}) = Q^2 \sum_{x=1}^{n} N_x(b_x - \bar{b})^2 + \left(\frac{n-1}{n}\right) \sum_{x=1}^{n} Qb_x[1 + Q(\phi_x - 1)]. \quad \text{(S8)}$$

The adjusted (net) raw *SSB* is obtained by subtracting the estimated random contribution, based on the sample variances:

$$\widehat{SSB}_{raw} = SSB_{raw} - \left(\frac{n-1}{n}\right) \sum_{x=1}^{n} s^2_{k_x}. \quad \text{(S9)}$$

Finally, the estimate of parametric *SSB* is obtained by rescaling the value in Equation S9 to account for sampling effects:

$$\widehat{SSB} = \widehat{SSB}_{raw}/Q^2. \quad \text{(S10)}$$

### LIFETIME REPRODUCTIVE SUCCESS

#### Parametric Sums of Squares

Under the assumption that an individual's reproduction in a given year does not affect its survival or reproductive output in any subsequent year, the expected *LRS* for individual *i* that dies at age *q* is the sum of its expected fecundity in years 1:*q*

$$E(k\bullet_{i,q}) = E(\overline{k\bullet}_q) = \sum_{x=1}^{q} E(k\bullet_{i,x}) = \sum_{x=1}^{q} b_x \ ,$$

and the expected variance in *LRS* for that individual is the sum of the expected variances in years 1:*q*

$$E(\sigma^2_{k\bullet,q}) = \sum_{x=1}^{q} \phi_x b_x.$$

Then,

$$E\left(SSE\bullet_q\right) = \sum_{i=1}^{D_q}\left(k\bullet_{i,q} - \overline{k\bullet}_q\right)^2 = D_q E(s_{k\bullet,q}^2) = D_q \sum_{x=1}^{q} \phi_x b_x \ ,$$

and the overall parametric within-groups sums of squares is

$$SSE\bullet_{parametric} = \sum_{q=1}^{n} D_q \sum_{x=1}^{q} \phi_x b_x. \qquad \text{(S11)}$$

Consider now the group-specific terms in parametric $SSB\bullet$. For individuals that die at age $q$, the relevant between-group term is $SSB\bullet_q = D_q(\overline{k\bullet}_q - \overline{k\bullet}.)^2$. Substituting for $E(\overline{k\bullet}_q) = \sum_{x=1}^{q} b_x$ and $\overline{k\bullet}. = 2$,

$$E\left(SSB\bullet_q\right) = D_q * \left(E\left(\overline{k\bullet}_q\right) - 2\right)^2 = D_q * \left((\sum_{x=1}^{q} b_x) - 2\right)^2,$$

so the overall parametric *SSB* becomes

$$SSE\bullet_{parametric} = \sum_{q=1}^{n} D_q * \left((\sum_{x=1}^{q} b_x) - 2\right)^2 \quad . \qquad \text{(S12)}$$

### ESTIMATION

**Case 1: Comprehensive Sampling**

Each group-specific term contributing to $\widehat{SSE\bullet}$ requires an estimate of the within-group variance $\sigma_{k\bullet,q}^2$. This estimator is calculated from the raw data as

$$\hat{\sigma}_{k\bullet,q}^2 = \frac{\sum_{i=1}^{D_q}(k\bullet_{i,x} - \overline{k\bullet}_q)^2}{D_q - 1} \ . \qquad \text{(S13)}$$

An estimator of the overall with-group sums of squares is then

$$\widehat{SSE\bullet} = \sum_{q=1}^{n} D_q \hat{\sigma}_{k\bullet,q}^2. \qquad \text{(S14)}$$

As with annual reproduction, the estimator for *SSB*• has to account for the fact that in empirical data, random variation in realized mean *LRS* for individuals that die at the same age *q* will increase $SSB\bullet_q$ above the parametric expectation. For comprehensive empirical data, the expected contribution to the between-group sums of squares from individuals that die at age *q* is

$$E\left(SSB\bullet_{q,comprehensive}\right) = E\left[D_q * \left(\overline{k\bullet}_q - 2\right)^2\right] = D_q * E\left[\left(\overline{k\bullet}_q - 2\right)^2\right].$$

As above, noting that $E\left[\left(\overline{k\bullet}_q - 2\right)^2\right] = \left[E(\overline{k\bullet}_q - 2)\right]^2 + var(\overline{k\bullet}_q - 2)$,

$$E\left(SSB\bullet_{q,comprehensive}\right) = D_q * \left[E(\overline{k\bullet}_q - 2)\right]^2 + D_q * var(\overline{k\bullet}_q - 2)$$

$$= D_q * var(\overline{k\bullet}_q) + D_q * \left((\textstyle\sum_{x=1}^{q} b_x) - 2\right)^2.$$

Now $var(\overline{k\bullet}_q) = var(k\bullet_{i,q})/D_q = \sigma^2_{k\bullet,q}/D_q$, so

$$E(SSB\bullet_{q,comprehensive}) = D_q * \sigma^2_{k\bullet,q}/D_q + D_q * \left((\textstyle\sum_{x=1}^{q} b_x) - 2\right)^2$$

$$= \sigma^2_{k\bullet,q} + D_q * \left((\textstyle\sum_{x=1}^{q} b_x) - 2\right)^2$$

$$= \textstyle\sum_{x=1}^{q} \phi_x b_x + D_q * \left((\sum_{x=1}^{q} b_x) - 2\right)^2$$

$$= E(SSE\bullet_q)/D_q + D_q * \left((\textstyle\sum_{x=1}^{q} b_x) - 2\right)^2 \quad ,$$

and the overall between-groups sums of squares is

$$E(SSB\bullet_{comprehensive}) = \frac{n-1}{n} \textstyle\sum_{q=1}^{n} \sum_{x=1}^{q} \phi_x b_x + \sum_{q=1}^{n} D_q * \left(\left(\sum_{x=1}^{q} b_x\right) - 2\right)^2$$

$$= \frac{n-1}{n} \textstyle\sum_{q=1}^{n} \hat{\sigma}^2_{k\bullet,q} + \sum_{q=1}^{n} D_q * \left(\left(\sum_{x=1}^{q} b_x\right) - 2\right)^2. \qquad \text{(S15)}$$

The quantity in Equation S15 exceeds the parametric expectation by the magnitude of the first term. Therefore, an unbiased estimator for $SSB\bullet$ is

$$\widehat{SSB\bullet} = SSB\bullet_{comprehensive} - \frac{n-1}{n} \textstyle\sum_{q=1}^{n} \hat{\sigma}^2_{k\bullet,q}. \qquad \text{(S16)}$$

After removing the random between-group contribution to $SSB\bullet_{comprehensive}$, the estimator $\widehat{SSB\bullet}$ accounts for the same two factors that contribute to parametric $SSB\bullet$: changes in fecundity with age, and differences in longevity. As shown above, these two factors can be partitioned as follows:

$$E(\widehat{SSB\bullet}_{longevity}) = \textstyle\sum_{q=1}^{n} D_q * \left(q\hat{\bar{b}} - 2\right)^2, \text{ and} \qquad \text{(S17)}$$

$$\widehat{SSB\bullet}_{between-age} = \widehat{SSB\bullet} - E(\widehat{SSB\bullet}_{longevity}). \qquad \text{(S18)}$$

**Case 2: Generalized Sampling Designs**

As before, we consider sampling at level $Q$ compared to comprehensive sampling and first develop an expectation for the raw sums of squares as a function of $Q$. We note that:

$E(\overline{k\bullet}_{q,raw}) = Q * E(\overline{k\bullet}_q) = Q \sum_{x=1}^{q} b_x$.
Target$\overline{k\bullet}_q = \sum_{x=1}^{q} b_x$ (for rescaling group-specific variances).
$\widehat{\Sigma b_x} = (\frac{1}{Q})\overline{k\bullet}_{q,raw}$
$E(\overline{k\bullet}_{.,raw}) = 2Q$

$s^2_{k\bullet,q}$ = sample variance of *LRS* among individuals that die at age $q$

$$E\left(s^2_{k\bullet,q}\right) = E(\overline{k\bullet}_{q,raw})\left[1 + \frac{E(\overline{k\bullet}_{q,raw})}{\sum_{x=1}^{q} b_x}\left(\frac{E(\sigma^2_{k\bullet,q})}{\sum_{x=1}^{q} b_x} - 1\right)\right]$$

$$= Q\sum_{x=1}^{q} b_x \left[1 + Q\left(\frac{\sum_{x=1}^{q}\phi_x b_x}{\sum_{x=1}^{q} b_x} - 1\right)\right].$$

Using results obtained above allows one to calculate the expected value of the raw, group-specific components of $SSE\bullet$ as

$$E\left(SSE\bullet_{q,raw}\right) = D_q * E\left(s^2_{k\bullet,q,raw}\right)$$

$$= D_q * Q\sum_{x=1}^{q} b_x \left[1 + Q\left(\frac{\sum_{x=1}^{q}\phi_x b_x}{\sum_{x=1}^{q} b_x} - 1\right)\right],$$

and to calculate the expected value of the overall within-group sums of squares as

$$E(SSE\bullet_{raw}) = \sum_{q=1}^{n} D_q * Q\sum_{x=1}^{q} b_x \left[1 + Q\left(\frac{\sum_{x=1}^{q}\phi_x b_x}{\sum_{x=1}^{q} b_x} - 1\right)\right]. \quad \text{(S19)}$$

Now we want to rescale the raw (empirical) variances to expected values under comprehensive sampling. An adjusted estimate of each within-group variance term is calculated as

$$\hat{\sigma}^2_{k\bullet,q,scaled} = \frac{\overline{k\bullet}_{q,raw}}{Q}\left[1 + \frac{1}{Q}\left(\frac{s^2_{k\bullet,q}}{\overline{k\bullet}_{q,raw}} - 1\right)\right],$$

which leads to an estimator of the within-group sums of squares as

$$\widehat{SSE\bullet}_q = D_q * \hat{\sigma}^2_{k\bullet,q,scaled} = D_q * \frac{\overline{k\bullet}_{q,raw}}{Q}\left[1 + \frac{1}{Q}\left(\frac{s^2_{k\bullet,q}}{\overline{k\bullet}_{q,raw}} - 1\right)\right],$$

and finally an estimator of the overall within-group sums of squares as

$$\widehat{SSE\bullet} = \sum_{q=1}^{n}\widehat{SSE\bullet}_q. \quad \text{(S20)}$$

For analysis of raw, empirical *LRS* data for between-group sums of squares, with analogy to Equation S15 for comprehensive sampling we can write

$$E\left(SSB\bullet_{q.raw}\right) = E\left(SSE\bullet_{q.raw}\right)/D_q + D_q * \left(E(\overline{k\bullet}_{q,raw}) - E(\overline{k\bullet}_{.,raw})\right)^2$$

$$= E\left(SSE\bullet_{q.raw}\right)/D_q + D_q * \left(Q(\textstyle\sum_{x=1}^{q} b_x) - 2Q\right)^2$$

$$= E\left(SSE\bullet_{q.raw}\right)/D_q + Q^2 D_q\left((\textstyle\sum_{x=1}^{q} b_x) - 2\right)^2.$$

Then the overall expectation for $SSB\bullet_{raw}$ is

$$E(SSB\bullet_{raw}) = \frac{n-1}{n}\sum_{q=1}^{n} E\big(SSE\bullet_{q.raw}\big)/D_q + Q^2\sum_{q=1}^{n} D_q\big((\sum_{x=1}^{q} b_x) - 2\big)^2, \qquad \text{(S21)}$$

and subtracting the expected contribution of random stochasticity to $raw\overline{k\bullet}_q$ produces this result

$$SSB\bullet_{raw,adjusted} = E(SSB\bullet_{raw}) - \frac{n-1}{n}\sum_{q=1}^{n} E\big(SSE\bullet_{q.raw}\big)/D_q$$

$$= Q^2\sum_{q=1}^{n} D_q\big((\sum_{x=1}^{q} b_x) - 2\big)^2. \qquad \text{(S22)}$$

The adjusted value in Equation S22 is then is rescaled to its expected value at comprehensive sampling:

$$\widehat{SSB}\bullet = SSB\bullet_{raw,adjusted}/Q^2. \qquad \text{(S23)}$$

Table S1. Variance partitioning analysis for seasonal reproduction by female black bears from Michigan. Assuming constant production of $N_1$ = 500 yearling females, these vital rates produce an adult female population size of 984 age 3+ individuals. Columns under "Raw data" show estimates from field samples reported by Waples et al. (2018b). Columns under "Scaled data" show variables that have been rescaled based on the estimated index of sampling intensity $Q$ = 0.097. The "Sums of squares" columns show age-specific within-age (*SSE*) and between-age (*SSB*) components. See Table 1 for notation, Table 3 for comparable data for males, and text for explanation of the calculations.

| | | | Raw data | | | | Scaled data | | | Sums of squares | | |
|---|---|---|---|---|---|---|---|---|---|---|---|---|
| Age | $l_x$ | $N_x$ | $\bar{k}_x$ | $s^2_{k_x}$ | $\phi_x$ | $B_x$ | $\hat{b}_x$ | $\hat{\sigma}^2_{k_{x,scaled}}$ | $\phi_x$ | $SSE$ | $SSE_{random}$ | $SSB_{raw}$ |
| 1 | 1 | 500 | 0.000 | | | | | | | | | |
| 2 | 0.767 | 384 | 0.000 | | | | | | | | | |
| 3 | 0.475 | 237 | 0.041 | 0.054 | 1.32 | 9.7 | 0.42 | 1.81 | 4.28 | 430.5 | 100.6 | 0.78 |
| 4 | 0.361 | 180 | 0.080 | 0.092 | 1.15 | 14.4 | 0.83 | 2.11 | 2.55 | 380.7 | 149.2 | 0.06 |
| 5 | 0.274 | 137 | 0.077 | 0.109 | 1.42 | 10.6 | 0.80 | 4.22 | 5.30 | 578.2 | 109.2 | 0.06 |
| 6 | 0.208 | 104 | 0.168 | 0.206 | 1.23 | 17.5 | 1.74 | 5.80 | 3.34 | 604.3 | 181.0 | 0.51 |
| 7 | 0.158 | 79 | 0.087 | 0.112 | 1.29 | 6.9 | 0.90 | 3.57 | 3.97 | 282.9 | 71.2 | 0.01 |
| 8 | 0.120 | 60 | 0.129 | 0.128 | 0.99 | 7.8 | 1.33 | 1.23 | 0.92 | 73.8 | 80.3 | 0.06 |
| 9 | 0.091 | 46 | 0.171 | 0.201 | 1.18 | 7.8 | 1.77 | 4.98 | 2.81 | 227.6 | 80.9 | 0.24 |
| 10 | 0.070 | 35 | 0.165 | 0.242 | 1.47 | 5.7 | 1.71 | 9.94 | 5.82 | 345.5 | 59.3 | 0.15 |
| 11+ | 0.053 | 105 | 0.155 | 0.191 | 1.23 | 16.3 | 1.60 | 5.41 | 3.37 | 567.7 | 168.3 | 0.34 |
| Totals | 3+ | 984 | | 1.33 | | 97 | | 39 | | 3491 | 1000 | 2.21 |

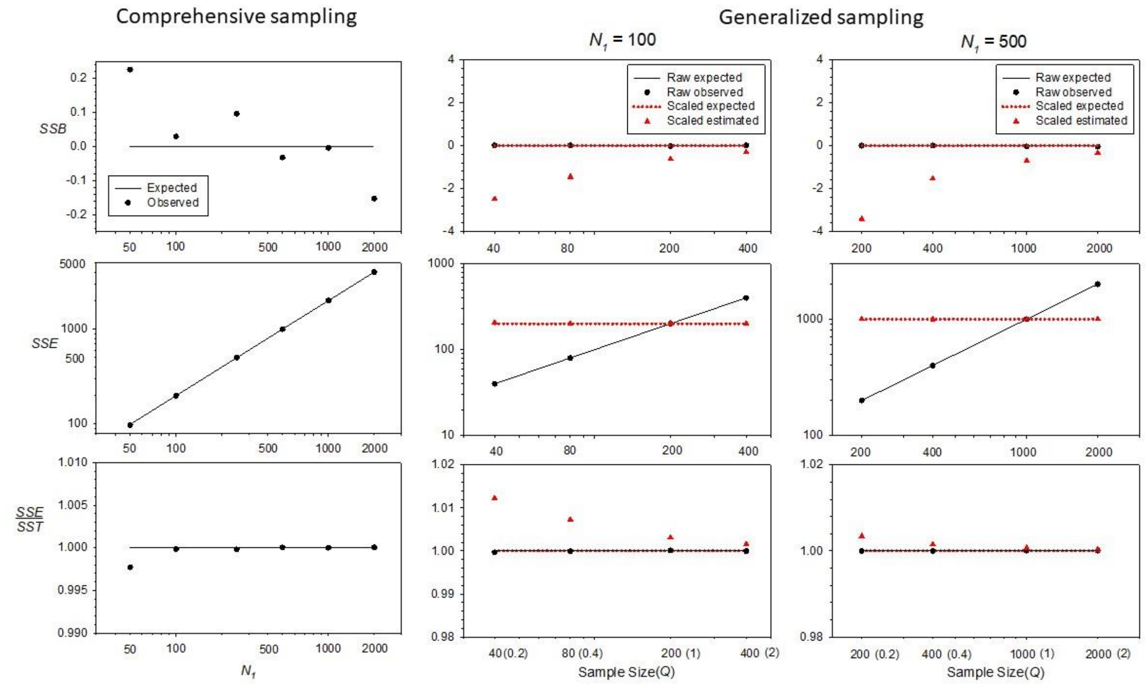


Figure S1. As in Figure 1 (main text) for annual reproduction, except these results are for a null model where fecundity was constant and variance in reproductive success within ages was Poisson (all $\phi_x = 1$).

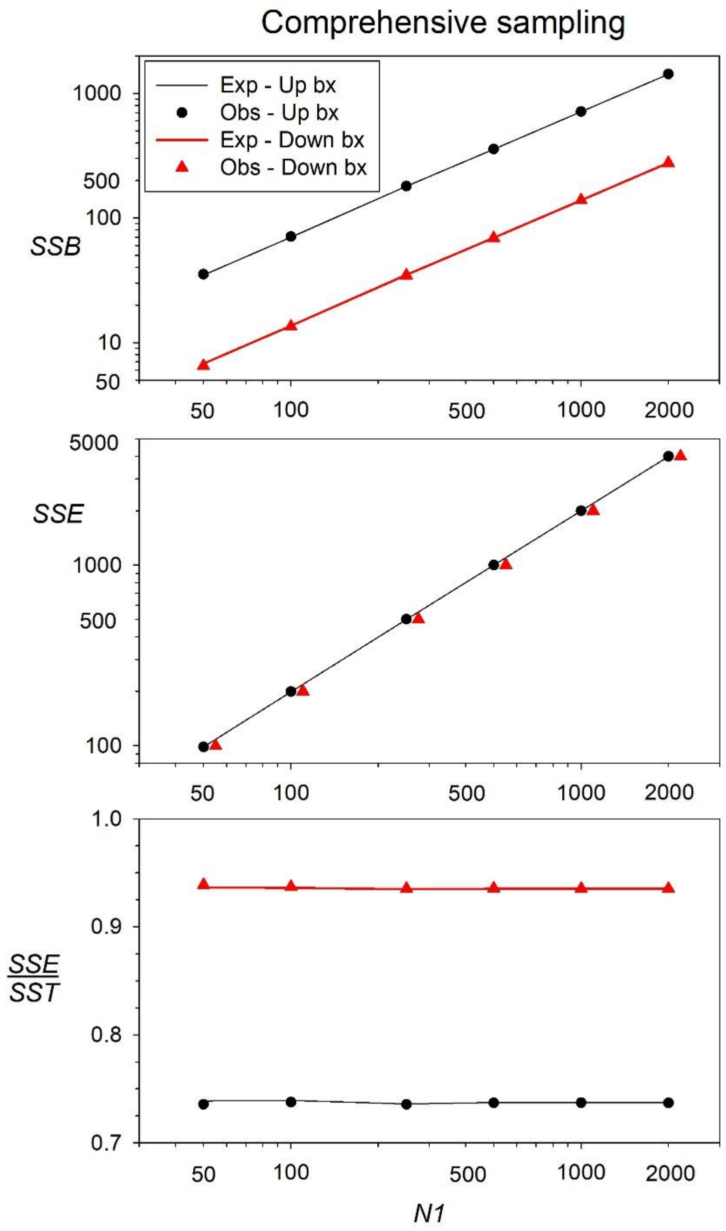


Figure S2. Comparison of expected ('exp', lines) and observed ('obs', symbols) sums of squares for scenarios with increasing fecundity with age ('Up bx', black) and decreasing fecundity ('Down bx', red). In all scenarios, sampling was comprehensive and $\phi = 1$. To facilitate the comparison, black lines and symbols repeat results shown in the left panel of Figure 1. In the middle panel for *SSE*, expectations for the two scenarios are essentially identical so only a single line is shown, and the red symbols for observed results for decreasing fecundity are jittered to the right.

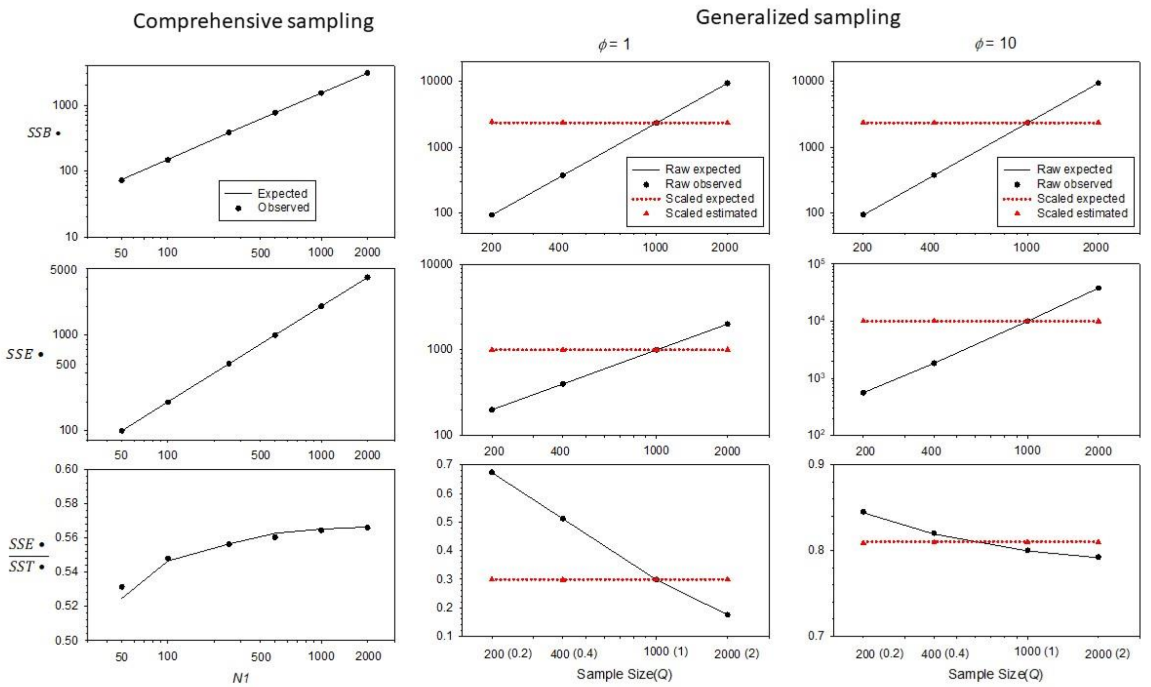


Figure S3. As in Figure 2 (main text) for lifetime reproduction, except these results are for a null model where fecundity was constant and variance in reproductive success within ages was Poisson (all $\phi_x = 1$).

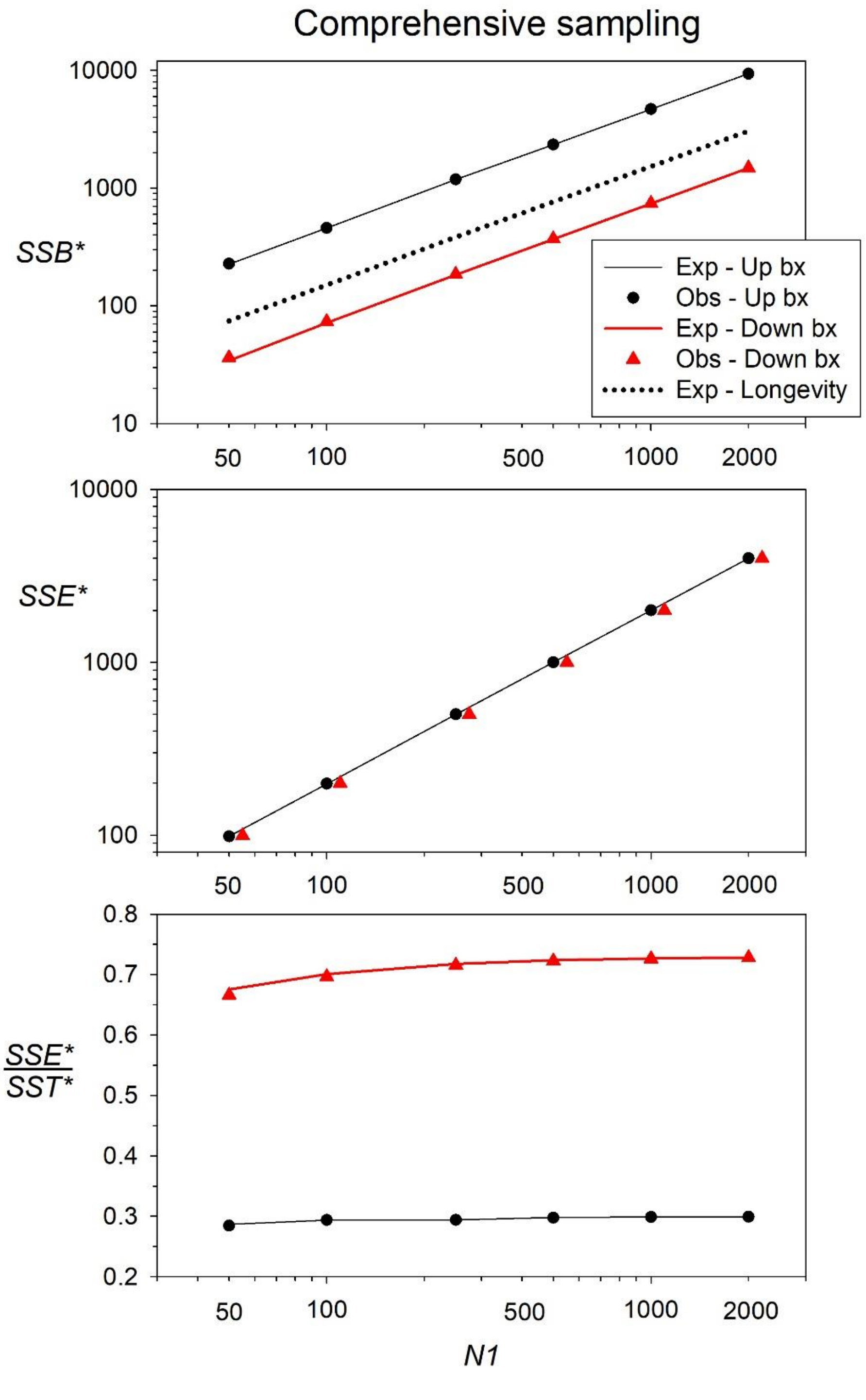


Figure S4. As in Figure S2, except for lifetime reproduction. In the top panel, the dotted line shows the expected value of *SSB*$\bullet_{Longevity}$.

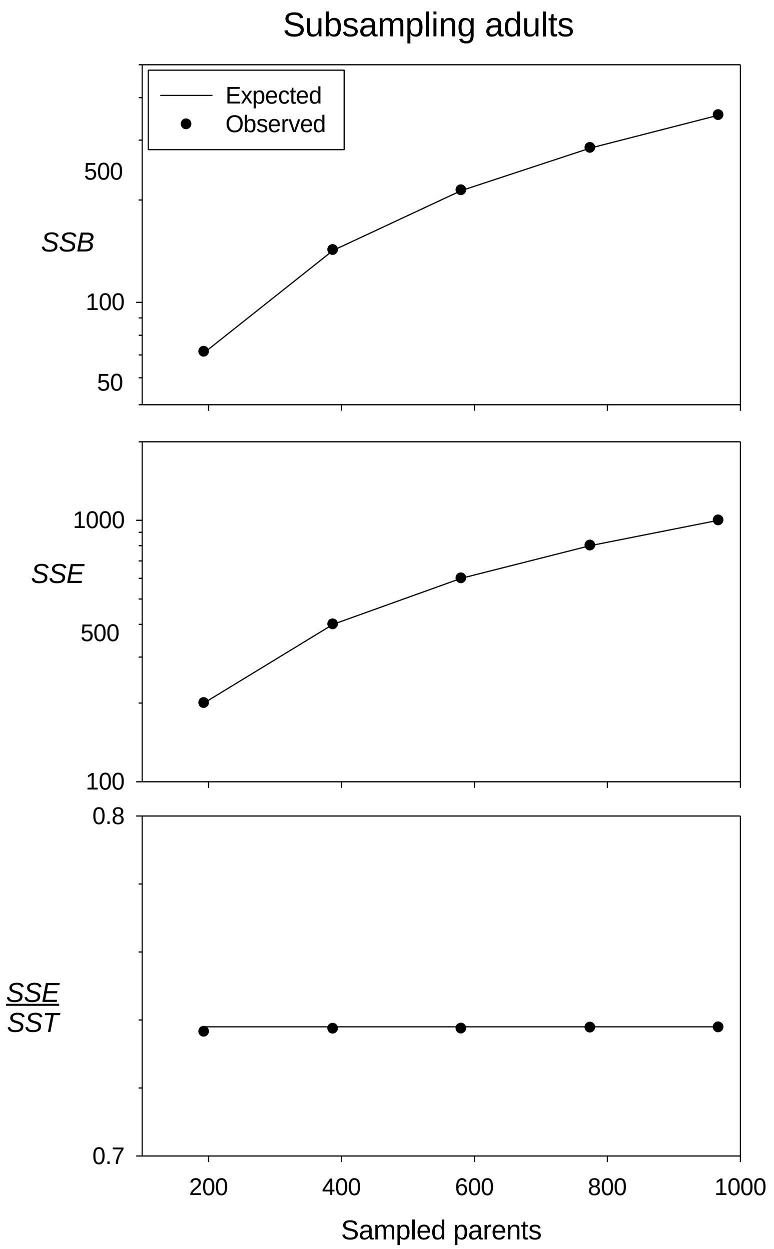


Figure S5. Expected and observed sums of squares for annual reproduction when subsampling adults. These simulations modeled the comprehensive sampling scenario in Figure 1, where fecundity increased with age, all $\phi$ = 1, and $N_1$ = 500, so the total number of adults was 967, results for which are shown in the rightmost datapoints. Other datapoints are for random subsamples of 20%, 40%, 60%, and 80% of the adults, with all of their assigned offspring. Note the log scale on the Y axes. The within- and between-age sums of squares both increase with the number of sampled parents, but their ratio is invariant.

**Box S1. Random and Deterministic Components to Annual *SSB***

For empirical data regarding annual reproduction, the overall between-groups sums of squares *SSB* has two components: a parametric one that reflects real differences in group means (changes in fecundity with age), and a random one that reflects stochastic sampling variation in group means. Even in the absence of parametric changes in fecundity with age, $E(SSB_{empirical})$ is greater than 0 because sample estimates of the group means differ by chance. As shown in the main text, the parametric component of *SSB* is

$$SSB_{parametric} = \sum_{x=1}^{n} N_x \, (b_x - \bar{b})^2 \quad .$$

For empirical data, the random component is based on the age-specific means in the samples ($\bar{k}_x$) and the weighted overall mean offspring number ($\bar{k}.$). The expectation for the random component depends on experimental design (esp sampling intensity), but for comprehensive sampling, each age-specific term is of the form

$$E(SSB_{x,random}) = s^2_{k_{i,x}},$$

where $s^2_{k_{i,x}}$ is the unbiased estimate of the variance in offspring number among individuals of age *x*. The overall $E(SSB_{random})$ across all ages is less than the sum of the age-specific terms, as variation in the group means is constrained by the overall weighted mean. With *n* age groups, there are *n*-1 degrees of freedom for $E(SSB_{random})$ rather than *n*, so the overall expectation is

$$E(SSB_{random}) = \left(\frac{n-1}{n}\right) \sum_{x=1}^{n} s^2_{k_{i,x}},$$

which is consistent with the fact that *SSB* has *n*-1 df in an ANOVA analysis.

Note that the random component to *SSB* in theory is independent of population or groups size (the vector of $N_x$ values). That this is also true in practice is illustrated in the figure below that reflects results of simulated data. These simulations used the 5-ages life table in Table B and considered a 16-fold range of $N_1$ values (60,120,240,480,960). Fecundity increased linearly with age, with $\phi$=1 for all ages, and sampling was comprehensive. Results shown are averaged across 100,000 replicate simulations. Plots 1 (dotted black line) and Plots 2 (black Xs) show expected and observed values, respectively, for $SSB_{random}$. Two points are evident from the plots: 1) realized $SSB_{random}$ actually is independent of *N*, as predicted; and 2) realized $SSB_{random}$ was consistently a bit higher than expected, by about 4-7%. However, even for small *N* the random component is expected to be only a small fraction of overall *SSB*. Furthermore, whereas the random component (and the absolute magnitude of the bias) is fixed, the parametric component of *SSB* increases linearly with *N*, so as population and group size increases, the small bias becomes increasingly less important (as indicated by the blue solid line). Even with $N_1$ as low as 60 (so group size for age 5 is only $N_5$ = 4), the upward bias to overall *SSB* is <1%, and it drops to <0.1% for larger population sizes.

Additional simulations were conducted to further examine patterns of bias in the random component of *SSB*. These simulations used comprehensive sampling and considered all combinations of constant *N* (all $N_x$ = 100) or decreasing *N* (with $N_1$ = 240, as above), and fecundity that was either constant, increasing, or decreasing with age. The table below shows the ratio (observed $SSB_{random}$)/(expected $SSB_{random}$), averaged across 100,000 replicates.

Figure 1.1. Plots 1 and 2 show observed and expected $SSB_{random}$ (left $Y$ axis) in simulations using a range of population sizes (indexed by $N_1$ on the $X$ axis; note log scale). These simulations had fecundity that increased with age and $\phi$=1 for all ages. Plot 3 (blue line) shows the bias [observed-expected] as a fraction of the overall parametric $SSB$ (with scale on the right $Y$ axis).

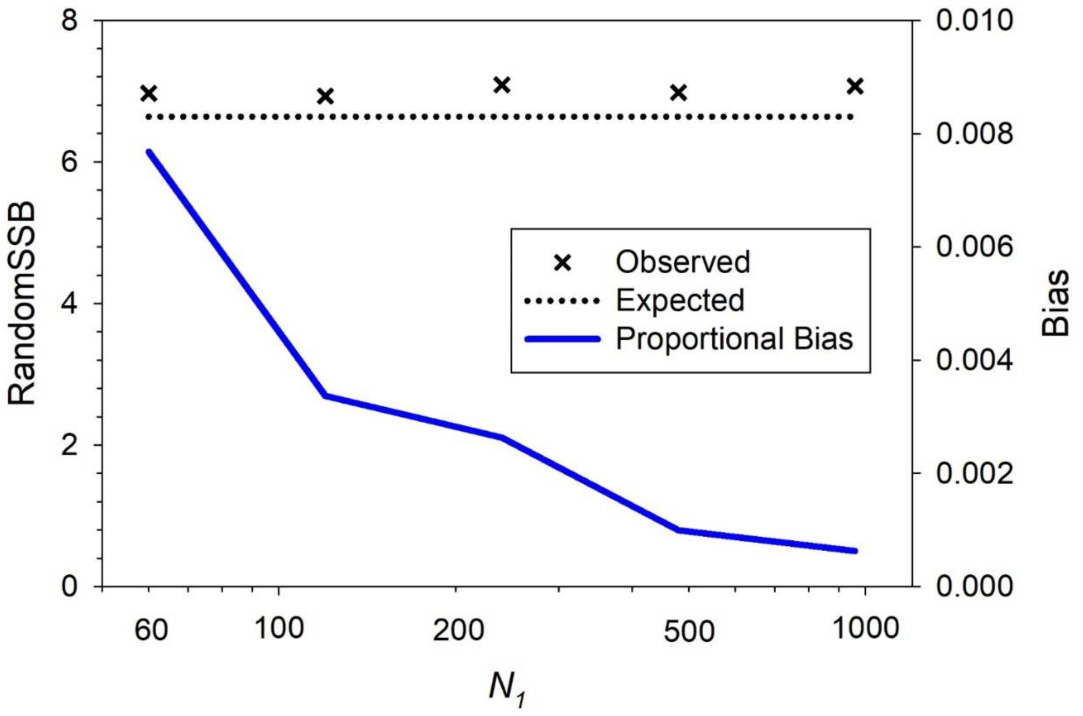


It is apparent that the pattern of age-specific fecundity controls the magnitude and direction of bias. When fecundity was constant, empirical $SSB$ was unbiased, regardless whether $N$ was constant or declining. When fecundity increased with age, bias was downwards with constant $N$ but upwards when group size declined with age. In the latter case, the correlation between $b_x$ and $N_x$ was -0.93. When fecundity declined with age, the pattern of bias was consistently down, and the magnitude of bias was greater when $N$ declined with age. In that case, the correlation between $b_x$ and $N_x$ was +0.93. Qualitatively similar results in all respects were found when simulating a 10-age life table with $N_x$ declining by 30% per year in the declining-$N$ scenario (data not shown).

Table 1.1. Values in the table are observed/expected $SSB_{random}$ for scenarios described in the text. Observed values are means across 100,000 replicate simulations.

| | | **Fecundity** | |
|---|---|---|---|
| **N** | **constant** | **up** | **down** |
| **constant** | 1.002 | 0.943 | 0.934 |
| **decreasing** | 1.000 | 1.033 | 0.872 |

**Box S2. Effects of age-specific fecundity on $SSB\bullet$ for lifetime reproduction**

This example illustrates calculation of lifetime $SSB\bullet$ for hypothetical populations with three different patterns of age-specific fecundity ($b_{x1}$, $b_{x2}$, $b_{x3}$), which have been scaled to values that will produce a stable population. Each population has the same number of individuals at each age ($N_x$) and the same number dying at each age ($D_q$), with both parameters reflecting 50% survival each year until age 5, after which time all survivors die. $\overline{k\bullet}_q = \sum_{x=1}^{q} b_x$ is cumulative mean *LRS* for individuals that died at age *q*, $\overline{k\bullet}.$ = mean *LRS* across all 500 individuals in the cohort is the weighted mean $\overline{k\bullet}_q$ and equals 2 for all stable populations, $SS\bullet_q = (\overline{k\bullet}_q - \overline{k\bullet}.)^2$ is the raw sums of squares for each age at death, and $SSB\bullet_q$ is $SS\bullet_q$ weighted by the group size ($D_q$). The totals are sums of $SS\bullet_q$ and $SSB\bullet_q$ for $q$=1-5.

| Age | $N_x$ | $D_q$ | $b_{x1}$ | $b_{x2}$ | $b_{x3}$ | $\overline{k\bullet}_{q,1}$ | $\overline{k\bullet}_{q,2}$ | $\overline{k\bullet}_{q,3}$ | $SS\bullet_{q,1}$ | $SS\bullet_{q,2}$ | $SS\bullet_{q,3}$ | $SSB\bullet_{q,1}$ | $SSB\bullet_{q,2}$ | $SSB\bullet_{q,3}$ |
|---|---|---|---|---|---|---|---|---|---|---|---|---|---|---|
| 1 | 500 | 250 | 0.56 | 1.03 | 1.24 | 0.56 | 1.03 | 1.24 | 2.1 | 0.9 | 0.6 | 517 | 233.7 | 144 |
| 2 | 250 | 125 | 1.12 | 1.03 | 0.99 | 1.69 | 2.07 | 2.23 | 0.1 | 0.0 | 0.1 | 12 | 1 | 7 |
| 3 | 125 | 63 | 1.69 | 1.03 | 0.74 | 3.37 | 3.10 | 2.98 | 1.9 | 1.2 | 1.0 | 119 | 76 | 60 |
| 4 | 62 | 31 | 2.25 | 1.03 | 0.50 | 5.62 | 4.13 | 3.47 | 13.1 | 4.5 | 2.2 | 407 | 141 | 67 |
| 5 | 31 | 31 | 2.81 | 1.03 | 0.25 | 8.44 | 5.17 | 3.72 | 41.4 | 10.0 | 3.0 | 1284 | 311 | 92 |
| | | | | | $\overline{k\bullet}.$ | 2.0 | 2.0 | 2.0 | | | | | | |
| | | | | | | | | Totals | 58.6 | 16.7 | 6.7 | 2339 | 762 | 370 |

The figure below shows these data graphically. Cumulative mean *LRS* ($\overline{k\bullet}_q$; Panel B) increases linearly with age at death when fecundity is constant (as in Population 2). If fecundity increases with age the increase in $\overline{k\bullet}_q$ becomes exponential (as in Population 1); if fecundity decreases with age the increase in $\overline{k\bullet}_q$ is asymptotic (as in Population 3). As a consequence of the much wider range of $\overline{k\bullet}_q$ values for Population 1 ($\overline{k\bullet}_5$ is 15 times as large as $\overline{k\bullet}_1$ compared to 3 times as large for Population 3), the squared deviations from the weighted mean [$SS\bullet_q = (\overline{k\bullet}_q - \overline{k\bullet}.)^2$] are also much larger for Population 1, as seen in Panel C, esp for older age at death. When the squared deviations are weighted by group size, producing group-specific values of $SSB\bullet_q$ (Panel D), the earlier ages at death are more heavily weighted, which reduces somewhat the disparity between Populations 1 and 3 ($\Sigma(SS\bullet)$ = 58.6/6.7 = 8.7 times as large for Population 1 compared to $\Sigma(SSB\bullet_q) = SSB\bullet$ = 2339/370 = 6.3 times as large for Population 1). Declining fecundity reduces overall $SSB\bullet$ (2339-370=1969 lower than with increasing fecundity) considerably more than does constant fecundity (1577 less), a result that presumably is related to the very compact range of group means ($\overline{k\bullet}_q$) for the declining-fecundity scenario.

In Population 2, constant $b_x$ eliminates any between-age fecundity differences, so $SSB\bullet$ can be attributed entirely to variation in age at death (longevity). This provides a useful reference point for evaluating the consequences of changing fecundity with age.

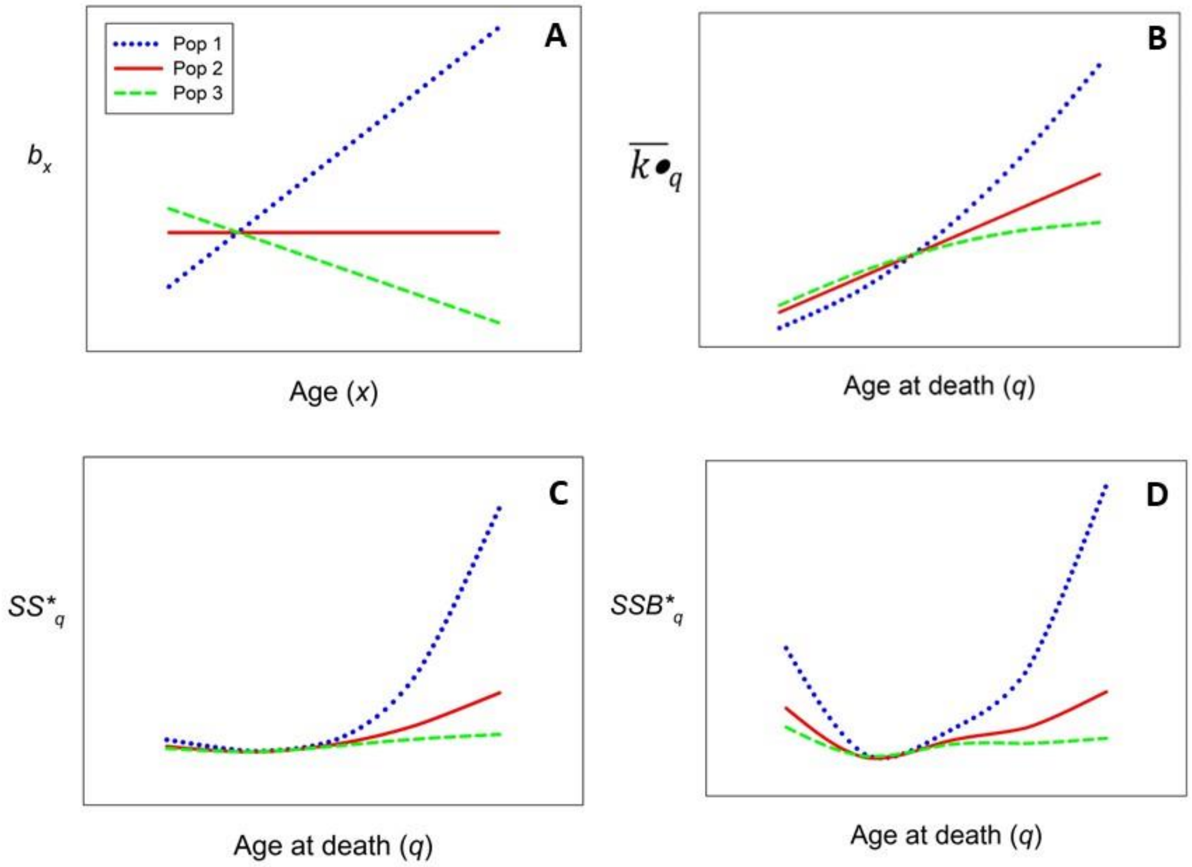
Pop 1
Pop 2
Pop 3
A
$b_x$
Age ($x$)
B
$\overline{k\bullet}_q$
Age at death ($q$)
C
$SS^*_q$
Age at death ($q$)
D
$SSB^*_q$
Age at death ($q$)

### Computer Code

Four sets of code are provided below: for comprehensive and generalized sampling for annual reproduction, and for comprehensive and generalized sampling for lifetime reproduction. All of these routines generate results that vary stochastically from run to run, unless controlled by a specific seed. With adequate replication, the mean results should resemble very closely those shown in the manuscript.

Annual reproduction, comprehensive sampling

```
NReps = 10000
baseN = c(500,250,125,62,31)
Ages = 1:length(baseN)
Multiple = c(0.1,0.2, 0.5, 1, 2,4) ## used to rescale population size
BigN = matrix(NA,length(Multiple),length(baseN))
for(k in 1:length(Multiple))  {
for(j in 1:length(baseN))  {
  BigN[k,j] = round(Multiple[k]*baseN[j])
  }}
TotN = rowSums(BigN)

##bx = rep(1,length(baseN)) ## constant fecundity
bx = 1:length(baseN)  ## fecundity increases with age
bx = rev(bx)
bxprime = bx*2*baseN[1]/sum(bx*baseN) ##stable N fecundity
phi = rep(1,length(baseN)) ##stable N phi
Vk = phi*bxprime
EV = sum(Vk)
OverallKbar = sum(bxprime*baseN)/sum(baseN)

Results = array(NA, dim = c(3,4,length(Multiple)))
rownames(Results) = c("Within","Between","Within/Total")
colnames(Results) = c("ParametricSS","Method1SS","ParametricVC","Method2VC")
dimnames(Results)[[3]] = Multiple*baseN[1]

MyMeans = array(NA, dim = c(NReps,length(baseN),length(Multiple)))
MyVars = MyMeans
ESSE = rep(NA, length(Multiple))
ESSB = ESSE
adjESSB = ESSE
EsigmaWithin = ESSE
EsigmaBetween = ESSE

rawSSB = matrix(NA,NReps,length(Multiple))
colnames(rawSSB) = Multiple
rawSSE = rawSSB
adjSSB = rawSSB

report = (NReps/10)*(1:10)  ### report progress every 10% of job

for (k in 1:length(Multiple))  {  ## for different population sizes

##Parametric sums of squares
  ESSE[k] = sum(BigN[k,]*Vk)
  ESSB[k] = EV*(length(baseN)-1)/length(baseN) + sum(BigN[k,]*(bxprime-OverallKbar)^2)
```

```
  adjESSB[k] = sum(BigN[k,]*(bxprime-OverallKbar)^2)
##Parametric variance components
   EsigmaWithin[k] = ESSE[k]/TotN[k]
   EsigmaBetween[k] = adjESSB[k]/TotN[k]

for (R in 1:NReps) {

if(R %in% report)  {
    print(paste0("Rep = ",R))
    flush.console()    }  ## end if

BigVector = 9999
for (j in 1:length(baseN))  {
if (phi[j] == 1) {kvector <- rpois(BigN[k,j], bxprime[j])}  ## use Poisson distribution for phi = 1 and negative binomial for phi > 1
  else { kvector <- rnbinom(BigN[k,j], mu = bxprime[j], size = bxprime[j]^2/(Vk[j]-bxprime[j])) }
BigVector = c(BigVector,kvector)
   MyMeans[R,j,k] = mean(kvector)
   MyVars[R,j,k]  = var(kvector) ## unbiased sample variance
   }  ## end for j

BigVector = BigVector[-1]
Kbar = sum(BigVector)/TotN[k]
rawSSE[R,k] = sum(BigN[k,]*MyVars[R,,k])
rawSSB[R,k] = sum(BigN[k,]*(MyMeans[R,,k]-Kbar)^2)  ## includes random variation in realized kbar
adjSSB[R,k] = rawSSB[R,k] - sum(MyVars[R,,k])*(length(baseN)-1)/length(baseN)

} ## end for R

} # end for k

Actualbx = t(colMeans(MyMeans))
colnames(Actualbx) = Ages
rownames(Actualbx) = Multiple*baseN[1]
ActualVk = t(colMeans(MyVars))
colnames(ActualVk) = Ages
rownames(ActualVk) = Multiple*baseN[1]
## Print these matrices if you want to see summaries of the simulated data
Actualbx
ActualVk
ActualVk/Actualbx

Results = matrix(NA,length(Multiple),8)  ## Method 1, estimating sums of squares
colnames(Results) = c("ESSE","SSE","ErawSSB","rawSSB","EadjSSB","adjSSB","EFractionWithin","FractionWithin")
rownames(Results) = Multiple*baseN[1]
Results[,1] = ESSE
Results[,2] = colMeans(rawSSE)
Results[,3] = ESSB
Results[,4] = colMeans(rawSSB)
Results[,5] =  adjESSB
Results[,6] =  colMeans(adjSSB)
Results[,7] =  Results[,1]/(Results[,1]+Results[,5])
Results[,8] =  Results[,2]/(Results[,2]+Results[,6])
Results
```

<u>Annual reproduction, generalized sampling</u>

```
## This code models a range of sampling efforts, indicated by the index Q
### Q < 1 represents proportional subsampling of a full cohort of offspring
## Q > 1 mimics sampling early life stages of a highly fecund species
## Q = 1 replicates comprehensive sampling
NReps = 10000
N = c(200,100,50,25,12)
#N = round(N/2)  # alternative ways of quickly rescaling population size without changing age structure
N = round(N*2.5)

Q = c(0.2,0.4,1,2)  ## index of sampling intensity
SampledOffspring = Q*2*N[1]
#bx = rep(1,length(N))
bx = 1:length(N)  ## increasing fecundity
##bx = rev(bx)  ## decreasing fecundity
bxprime = bx*2*N[1]/sum(bx*N)  ## rescales bx to values for a stable population
phi = rep(1,length(N)) ##stable N phi

sampleVk = matrix(NA,length(Q),length(N))
samplebx = sampleVk
for (q in 1:length(Q)) {
for (j in 1:length(N)) {
  samplebx[q,j] = bxprime[j]*Q[q]
  sampleVk[q,j] = samplebx[q,j]*(1 + (samplebx[q,j]/bxprime[j])*(phi[j]-1))  ## rescaling Vk per Crow and Morton 1955
  }}

SampleKbar = 1:length(Q)
for (q in 1:length(Q))  {
  SampleKbar[q] = sum(samplebx[q,]*N)/sum(N)  }

EV = matrix(NA,length(Q),length(N))
ESSE = 1:length(Q)
ESSB = 1:length(Q)
adjESSB = 1:length(Q)
for (q in 1:length(Q)) {
for (j in 1:length(N)) {
  EV[q,j] = sampleVk[q,j]
  }}

##Parametric sums of squares
for (q in 1:length(Q)) {
  ESSE[q] = sum(N*EV[q,])
  ESSB[q] = sum(EV[q,])* (length(N)-1)/length(N) + sum(N*(samplebx[q,]-SampleKbar[q])^2)
  adjESSB[q] = sum(N*(samplebx[q,]-SampleKbar[q])^2)  }

MyMeans = array(NA, dim = c(NReps,length(N),length(Q)))
MyVars = MyMeans
rawSSB = matrix(NA,NReps,length(Q))
colnames(rawSSB) = Q
rawSSE = rawSSB
SSBprime = rawSSB
scaledSSE = rawSSB
scaledSSBprime1 = rawSSB  ## adjust, then rescale
```

```
scaledSSBprime2 = rawSSB  ## rescale, then adjust
scaledSSB = rawSSB
EDeterministic = 0

report = (NReps/10)*(1:10)  ### report progress every 10% of job

for (R in 1:NReps) {

if(R %in% report)  {
    print(paste0("Rep = ",R))
    flush.console()    }  ## end if

q = length(Q)  ## first simulate maximum number of offspring

BigOff = rep(NA,2)
bxhat = rep(NA, length(N))
Vkhat = bxhat

for (j in 1:length(N))  {
ParentAge = rep(j,N[j])
if (phi[j] == 1) {kvector <- rpois(N[j], samplebx[q,j])}
  else { kvector <- rnbinom(N[j], mu = samplebx[q,j], size = samplebx[q,j]^2/(sampleVk[q,j]-samplebx[q,j])) }
Add = cbind(kvector,ParentAge)
BigOff = rbind(BigOff,Add)
   MyMeans[R,j,q] = mean(kvector)
   bxhat[j] = mean(kvector)/Q[q]
   MyVars[R,j,q]  = var(kvector) ## unbiased sample variance
   if(mean(kvector)>0)  {Vkhat[j] = bxhat[j]*(1 + (1/Q[q])*(var(kvector)/mean(kvector) - 1)) } else {Vkhat[j] = 0}
      }  ## end for j

Vkhat[Vkhat<0] <- 0
BigOff = BigOff[-1,]
Kbar = sum(BigOff[,1])/sum(N)
scaledKbar = Kbar/Q[q]
rawSSE[R,q] = sum(N*MyVars[R,,q])
rawSSB[R,q] = sum(N*(MyMeans[R,,q]-Kbar)^2)
SSBprime[R,q] = rawSSB[R,q] - sum(MyVars[R,,q])*(length(N)-1)/length(N)
scaledSSBprime1[R,q] = SSBprime[R,q]/Q[q]^2
scaledSSE[R,q] = sum(N*Vkhat)
scaledSSB[R,q] = sum(N*(bxhat-scaledKbar)^2)
scaledSSBprime2[R,q] = scaledSSB[R,q] - sum(Vkhat)*(length(N)-1)/length(N)

### Now do subsampling
### First, eliminate parents that produced no offspring
YesOff = subset(BigOff,BigOff[,"kvector"]>0)
### Now, convert the matrix of parents to a matrix of offspring
TotOff = sum(BigOff[,"kvector"])
MyOffspring = rep(NA,2)
for(k in 1:nrow(YesOff))  {
nOff = YesOff[k,1]
Moms = rep(k,nOff)
MomAge = rep(YesOff[k,2],nOff)
chunk = cbind(Moms,MomAge)
```

```
MyOffspring = rbind(MyOffspring,chunk)
} ## end for k
MyOffspring = MyOffspring[-1,]
##head(MyOffspring)
##table(MyOffspring[,"MomAge"])

for (q in 1:(length(Q)-1) ) {
LessOffspring = MyOffspring[sample(nrow(MyOffspring),SampledOffspring[q],replace=FALSE),]

bxhat = rep(NA, length(N))
Vkhat = bxhat

for (j in 1:length(N))  {
  SomeAges = subset(LessOffspring, LessOffspring[,"MomAge"] == j)
  MyMeans[R,j,q] = nrow(SomeAges)/N[j]
  bxhat[j] = MyMeans[R,j,q]/Q[q]
  skew = table(SomeAges[,"Moms"])
  SS = sum(skew^2)
  V = SS/N[j] - MyMeans[R,j,q]^2  ## population variance
  MyVars[R,j,q]  = V*N[j]/(N[j]-1) ## unbiased sample variance
  if(MyMeans[R,j,q]>0)  {Vkhat[j] = bxhat[j]*(1 + (1/Q[q])*(MyVars[R,j,q]/MyMeans[R,j,q] - 1)) } else {Vkhat[j] = 0}
    }  ## end for j
Vkhat[Vkhat<0] <- 0

Kbar = SampledOffspring[q]/sum(N)
scaledKbar = Kbar/Q[q]
rawSSE[R,q] = sum(N*MyVars[R,,q])
rawSSB[R,q] = sum(N*(MyMeans[R,,q]-Kbar)^2)
SSBprime[R,q] = rawSSB[R,q] - sum(MyVars[R,,q])*(length(N)-1)/length(N)
scaledSSBprime1[R,q] = SSBprime[R,q]/Q[q]^2
scaledSSE[R,q] = sum(N*Vkhat)
scaledSSB[R,q] = sum(N*(bxhat-scaledKbar)^2)
scaledSSBprime2[R,q] = scaledSSB[R,q] - sum(Vkhat)*(length(N)-1)/length(N)

} # end for q

} ## end for R

Actualbx = colMeans(MyMeans)
colnames(Actualbx) = Q
rownames(Actualbx) = N
ActualVk = colMeans(MyVars)
colnames(ActualVk) = Q
rownames(ActualVk) = N
## Print these matrices if you want to see summaries of the simulated data
Actualbx
ActualVk
ActualVk/Actualbx

Results = matrix(NA,length(Q),8)
colnames(Results) = c("ESSE","SSE","ErawSSB","rawSSB","ESSBprime","SSBprime","EFractionWithin","FractionWithin")
rownames(Results) = Q
Results[,1] = ESSE
```

```
Results[,2] = colMeans(rawSSE)
Results[,3] = ESSB
Results[,4] = colMeans(rawSSB)
Results[,5] =  adjESSB
Results[,6] =  colMeans(SSBprime)
Results[,7] =  Results[,1]/(Results[,1]+Results[,5])
Results[,8] =  Results[,2]/(Results[,2]+Results[,6])

ResultsB = matrix(NA,length(Q),9)
colnames(ResultsB) =
c("ESSE","ScaledSSE","EScaledSSB","ScaledSSB","ESSBprime","ScaledSSBprime1","ScaledSSBprime2","EFractionWithin","FractionWithin")
rownames(ResultsB) = Q
ResultsB[,1] = ESSE[3]
ResultsB[,2] = colMeans(scaledSSE,na.rm=T)
ResultsB[,3] = ESSB[3]
ResultsB[,4] = colMeans(scaledSSB,na.rm=T)
ResultsB[,5] =  adjESSB[3]
ResultsB[,6] =  colMeans(scaledSSBprime1,na.rm=T)
ResultsB[,7] =  colMeans(scaledSSBprime2,na.rm=T)
ResultsB[,8] =  ResultsB[,1]/(ResultsB[,1]+ResultsB[,5])
ResultsB[,9] =  ResultsB[,2]/(ResultsB[,2]+ResultsB[,6])

Results  ## compares raw data with expected values based on values of Q
ResultsB ## rescales sums of squares and compares with parametric expectations under comprehensive sampling
##scaledSSBprime1 is the method used in the main text; it subtracts the expected random component from the raw SSB, then rescales the result
##scaledSSBprime2 is an alternative method mentioned in the SI; it first rescales the age-specific sample variances, then subtracts the expected random component from the rescaled SSB.
```

<u>Lifetime reproduction, comprehensive sampling</u>

```
##This code models 'parametric' variance components under comprehensive sampling, where the full 2*N1 offspring in a cohort are sampled
NReps = 10000
baseN = c(500,250,125,62,31)
AL = length(baseN)
alpha = 1
Ages = 1:length(baseN)
Multiple = c(0.1,0.2, 0.5, 1, 2,4) ## used to rescale population size
BigN = matrix(NA,length(Multiple),length(baseN))
for(k in 1:length(Multiple))  {
for(j in 1:length(baseN))  {
  BigN[k,j] = round(Multiple[k]*baseN[j])
  }}

##bx = rep(1,length(baseN)) ## constant fecundity
bx = 1:length(baseN)  ## fecundity increases with age
##bx = rev(bx)
bxprime = bx*2*baseN[1]/sum(bx*baseN) ##stable N fecundity
phi = rep(1,length(baseN)) ##stable N phi
Vk = phi*bxprime
EV = sum(Vk)
```

```
Ekbar = 2*baseN[1]/sum(baseN)

MyMeans = array(NA, dim = c(NReps,length(baseN),length(Multiple)))
MyVars = MyMeans
BigLRO = matrix(NA,NReps,length(Multiple))
Bigkbar = BigLRO
ESSE = rep(NA, length(Multiple))
ESSB = ESSE
adjESSB = ESSE

rawSSB = matrix(NA,NReps,length(Multiple))
colnames(rawSSB) = Multiple
rawSSE = rawSSB
adjSSB = rawSSB

ELong = 1:length(Multiple)

report = (NReps/10)*(1:10)  ### report progress every 10% of job

for (M in 1:length(Multiple))  {

Died = rep(NA,AL)
Nx = BigN[M,]
for (q in 1:(AL-1))  { Died[q] = Nx[q] - Nx[q+1] }
Died[AL] = Nx[AL]
NAdults = sum(Nx)
cohort = Nx[1] ## total number in the cohort

ELong[M] = sum(Died*(Ages*Ekbar-2)^2)

within = rep(NA,length(baseN))
between = within
bit = within
for (q in 1:length(baseN)) {
  within[q] = Died[q]*sum(bxprime[1:q]*phi[1:q])
  bit[q] = sum(bxprime[1:q]*phi[1:q])
  between[q] = bit[q] + Died[q]*(sum(bxprime[1:q])-2)^2
  }  # end for q
ESSE[M] = sum(within)
ESSB[M] = sum(between)
adjESSB[M] = ESSB[M] - sum(bit)

#########
for (R in 1:NReps)  {

if(R %in% report)  {
   print(paste0("multiple = ",M, " Replicate = ",R))
   flush.console()    }  ## end if

Pdata = matrix(NA,cohort,AL)
IDs = 1:cohort
Pdata = cbind(IDs,Pdata)
```

```
## Model reproduction at age 1
if (phi[1] == 1) {Offspring <- rpois(cohort,bxprime[1])} else {
Offspring <- rnbinom(cohort, mu = bxprime[1], size = bxprime[1]^2/(Vk[1]-bxprime[1])) }
Pdata[,2] = Offspring

Survivors = list()
for (q in 1:AL)  {
 Survivors[[q]] = 1:Nx[q]  }

Survivors[[1]] = IDs
## get survivors to subsequent ages , and their offspring production
for (j in 2:AL)  {
  NewN = Nx[j]
  Survivors[[j]] = sample(Survivors[[j-1]],NewN,replace=F)
  if (phi[j] == 1) {NewOffspring <- rpois(NewN,bxprime[j])} else {
     NewOffspring <- rnbinom(cohort, mu = bxprime[j], size = bxprime[j]^2/(Vk[j]-bxprime[j])) }
  for (k in 1:NewN) {
      row = Survivors[[j]][k]
      Pdata[row,j+1] = NewOffspring[k] }  ## end for k
    } ## end for j

V = 1:AL
for (q in 1:AL)  {
V[q] = var(Pdata[,q+1],na.rm=T)  ## variance in offspring number among survivors at each age
}  # end for q

LRO = rowSums(Pdata[,-1],na.rm=T)  ## lifetime number of offspring for each member of the cohort
BigLRO[R,M] = var(LRO)
Bigkbar[R,M] = mean(LRO)

              ##Get age at death for each member of the cohort
              PD = Pdata[,-1]
              AgeDeath = rep(AL,cohort)
              for (k in 1:cohort)  {
              X = which(is.na(PD[k,]))
              if(length(X)>0)  {AgeDeath[k] = X[1]-1}
              }  # end for k
              ##table(AgeDeath)

Pdata2 = cbind(Pdata,LRO,AgeDeath)

for (j in 1:length(baseN))  {
 group = subset(Pdata2,Pdata2[,"AgeDeath"] == j)
 MyMeans[R,j,M] = mean(group[,"LRO"])
 MyVars[R,j,M] = var(group[,"LRO"])
 } ## end for j

 rawSSE[R,M] = sum(Died*MyVars[R,,M])
 rawSSB[R,M] = sum(Died*(MyMeans[R,,M]-mean(LRO))^2)
 adjSSB[R,M] =  rawSSB[R,M] - sum(MyVars[R,,M])

}  # end for R
}  # end for M
```

```
BigSSE = colMeans(rawSSE)
BigSSB = colMeans(rawSSB)
SST = BigSSE+BigSSB
BigAdjSSB = colMeans(adjSSB)
FSSE = BigSSE/SST
EFSSE = ESSE/(ESSE+ESSB)

ResultsLRS = cbind(BigSSE,ESSE,BigSSB,ESSB,BigAdjSSB,adjESSB,FSSE,EFSSE)
rownames(ResultsLRS) = Multiple*baseN[1]
ResultsLRS
```

<u>Lifetime reproduction, generalized sampling</u>

```
NReps = 10000
N = c(500,250,125,62,31)
#N = c(100,50,25,12,6)
AL = length(N)
Died = rep(NA,AL)
for (q in 1:(AL-1))  { Died[q] = N[q] - N[q+1] }
Died[AL] = N[AL]
NAdults = sum(N)
cohort = N[1] ## total number in the cohort

alpha = 1
Ages = 1:length(N)
Q = c(0.2,0.4,1,2)  ## index of sampling intensity

bx = rep(1,length(N)) ## constant fecundity
bx = 1:length(N)  ## fecundity increases with age
bxprime = bx*2*N[1]/sum(bx*N) ##stable N fecundity
phi = rep(1,length(N)) ##stable N phi
Vk = phi*bxprime
EV = sum(Vk)

sampleVk = matrix(NA,length(Q),length(N))
samplebx = sampleVk
for (k in 1:length(Q)) {
for (j in 1:length(N)) {
  samplebx[k,j] = bxprime[j]*Q[k]
  sampleVk[k,j] = samplebx[k,j]*(1 + (samplebx[k,j]/bxprime[j])*(phi[j]-1))  ## rescaling Vk per Crow and Morton 1955
  }}

MyMeans = array(NA, dim = c(NReps,length(N),length(Q)))
MyVars = MyMeans
ScaledMeans = MyMeans
ScaledVars = MyMeans
BigLRO = matrix(NA,NReps,length(Q))
Bigkbar = BigLRO
ESSE = rep(NA, length(Q))
ESSB = ESSE
adjESSB = ESSE

rawSSB = matrix(NA,NReps,length(Q))
```

```
colnames(rawSSB) = Q
rawSSE = rawSSB
adjSSB = rawSSB
scaledSSB = rawSSB
scaledSSE = rawSSB
scaledSSBprime1 = rawSSB
scaledSSBprime2 = rawSSB

report = (NReps/10)*(1:10)  ### report progress every 10% of job

for (k in 1:length(Q))  {

within = rep(NA,length(N))
between1 = within
between2 = within
for (q in 1:length(N)) {
  between1[q] = Q[k]*sum(bxprime[1:q])*(1+Q[k]*(sum(bxprime[1:q]*phi[1:q])/sum(bxprime[1:q]) - 1))
  within[q] = between1[q]*Died[q]
  between2[q] = Died[q]*Q[k]^2*(sum(bxprime[1:q])-2)^2
  }  # end for q
ESSE[k] = sum(within)
ESSB[k] = sum(between2) + ((AL-1)/AL)*sum(between1)
adjESSB[k] = sum(between2)

Wbit = rep(NA,length(N))
Bbit = Wbit
for (q in 1:length(N)) {
  Wbit[q] = sum(bxprime[1:q]*phi[1:q])
  Bbit[q] = (sum(bxprime[1:q])-2)^2
  }  ## end for q
EscaledSSE = sum(Died*Wbit)
EscaledSSB = sum(Died*Bbit)

##############################################
for (R in 1:NReps)  {

if(R %in% report)  {
    print(paste0("Q = ",k, " Replicate = ",R))
    flush.console()    }  ## end if

Pdata = matrix(NA,cohort,AL)
IDs = 1:cohort
Pdata = cbind(IDs,Pdata)

## Model reproduction at age 1
if (phi[1] == 1) {Offspring <- rpois(cohort,samplebx[k,1])} else {
Offspring <- rnbinom(cohort, mu = samplebx[k,1], size = samplebx[k,1]^2/(sampleVk[k,1]-samplebx[k,1])) }
Pdata[,2] = Offspring

Survivors = list()
for (q in 1:AL)  {
 Survivors[[q]] = 1:N[q]  }
```

```
Survivors[[1]] = IDs
## get survivors to subsequent ages , and their offspring production
for (j in 2:AL)  {
  NewN = N[j]
  Survivors[[j]] = sample(Survivors[[j-1]],NewN,replace=F)
if (phi[j] == 1) {NewOffspring <- rpois(cohort,samplebx[k,j])} else {
  NewOffspring <- rnbinom(cohort, mu = samplebx[k,j], size = samplebx[k,j]^2/(sampleVk[k,j]-samplebx[k,j])) }
  for (kk in 1:NewN) {
      row = Survivors[[j]][kk]
      Pdata[row,j+1] = NewOffspring[kk] }  ## end for kk
    } ## end for j

V = 1:AL
for (q in 1:AL)  {
V[q] = var(Pdata[,q+1],na.rm=T)  ## variance in offspring number among survivors at each age
}  # end for q

LRO = rowSums(Pdata[,-1],na.rm=T)  ## lifetime number of offspring for each member of the cohort
BigLRO[R,k] = var(LRO)
Bigkbar[R,k] = mean(LRO)

               ##Get age at death for each member of the cohort
               PD = Pdata[,-1]
               AgeDeath = rep(AL,cohort)
               for (kk in 1:cohort)  {
                 X = which(is.na(PD[kk,]))
                 if(length(X)>0)  {AgeDeath[kk] = X[1]-1}
                 }  # end for kk
               ##table(AgeDeath)

Pdata2 = cbind(Pdata,LRO,AgeDeath)

for (j in 1:length(N))  {
 group = subset(Pdata2,Pdata2[,"AgeDeath"] == j)
 MyMeans[R,j,k] = mean(group[,"LRO"])
 MyVars[R,j,k] = var(group[,"LRO"])
 ScaledMeans[R,j,k] = MyMeans[R,j,k]/Q[k]
 if(ScaledMeans[R,j,k]>0) {ScaledVars[R,j,k] = ScaledMeans[R,j,k]*(1 + (1/Q[k])*(MyVars[R,j,k]/MyMeans[R,j,k]-1)) } else
{ScaledVars[R,j,k]=0}
 } ## end for j

ScaledMeans[R,,k][ScaledMeans[R,,k]<0] <- 0
Kbar = sum(LRO)/sum(Died)
scaledKbar = Kbar/Q[k]

 rawSSE[R,k] = sum(Died*MyVars[R,,k])
 rawSSB[R,k] = sum(Died*(MyMeans[R,,k]-mean(LRO))^2)
 adjSSB[R,k] =  rawSSB[R,k] - ((AL-1)/AL)*sum(MyVars[R,,k])
 scaledSSE[R,k] = sum(Died*ScaledVars[R,,k])
 scaledSSB[R,k] = sum(Died*(ScaledMeans[R,,k]-scaledKbar)^2)
 scaledSSBprime1[R,k] = adjSSB[R,k]/Q[k]^2
 scaledSSBprime2[R,k] =  scaledSSB[R,k] -  ((AL-1)/AL)*sum(ScaledVars[R,,k])
```

```
}  # end for R

}  # end for M

BigSSE = colMeans(rawSSE)
BigSSB = colMeans(rawSSB)
SST = BigSSE+BigSSB
BigAdjSSB = colMeans(adjSSB)
FSSE = BigSSE/SST
EFSSE = ESSE/(ESSE+ESSB)

BigscaledSSE = colMeans(scaledSSE,na.rm=T)
BigscaledSSB1 = colMeans(scaledSSBprime1,na.rm=T)
BigscaledSSB2 = colMeans(scaledSSBprime2,na.rm=T)
scaledFSSE = BigscaledSSE/(BigscaledSSE+BigscaledSSB1)
EscaledFSSE = EscaledSSE/(EscaledSSE+EscaledSSB)

rawResultsLRS = cbind(ESSE,BigSSE,ESSB,BigSSB,adjESSB,BigAdjSSB,FSSE,EFSSE)
scaledResultsLRS = cbind(EscaledSSE,BigscaledSSE,EscaledSSB,BigscaledSSB1,BigscaledSSB2,scaledFSSE,EscaledFSSE)
rawResultsLRS  ## compares raw data with expected values based on values of Q
scaledResultsLRS ## rescales sums of squares and compares with parametric expectations under comprehensive sampling
##BigscaledSSB1 is the method used in the main text; it subtracts the expected random component from the raw SSB*, then rescales the result
##BigscaleddSSB2 is an alternative method mentioned in the SI; it first rescales the age-specific sample variances, then subtracts the expected random component from the rescaled SSB*.

rawResultsLRS
scaledResultsLRS
```
